# Axonal theta oscillations evoke bursting in target hippocampal subregions

**DOI:** 10.64898/2026.01.21.700016

**Authors:** Samuel B. Lassers, Gregory J. Brewer

## Abstract

Local field potentials (LFPs) measured in the extracellular matrix of the brain are postulated to arise from the integration of synaptic ionic currents and spread by volume conduction. However, there is a lack of consensus on whether these spatiotemporal voltage gradients are just an epiphenomenon of spiking or if the LFPs play a functional role in information processing. To examine a potential functional role of LFPs in information processing, we developed a microfluidic device that allows neurons from the hippocampal formation to self-wire through microfluidic channels, effectively isolating the activity of single axons between subregions of the network. We recorded spontaneous theta-band activity (4-10 Hz) in these axons whose power spectra were independent of simultaneous spiking activity. A sparse set of axons from the CA3 into the CA1 had the highest theta amplitudes. Source neurons for the axonal theta were identified through cross correlation. Functionally, sparse axonal theta phase and amplitude correlated with target subregional spiking and more strongly with burst length. These results suggest that theta voltage oscillations in axons may contribute to activation of slow voltage-gated calcium channels to drive stronger synaptic release of transmitter to coordinate hippocampal activity between subregions. We propose that theta oscillations are controlled by specific ion channels distinct from those that generate spikes, a multiplex coding mechanism for inter-regional communication with implications for routing, executive control, disease states and artificial neural networks.

## Introduction

Technically, every local transmembrane current in the brain contributes to an extracellular voltage commonly referred to as a local field potential (LFP)^1^. From the first recordings of EEG voltage oscillations on the scalp recounted by^2^ to modern depth electrodes in animals and humans, common frequencies were found to correspond to certain motor behaviors and problem solving (beta, 13-30 Hz), relaxation (alpha, 8-13 Hz), sleep (delta, 1-4 Hz; spindles, 10-16 Hz), and planning, learning and memory (theta 4-10 Hz). The source of these LFP’s remains uncertain but could be important for interregional communication since synchronized oscillations are observed and interpreted as evidence for communication between brain subregions^3^. Many assume they are the result of summation of simultaneous synaptic currents (EPSC’s), thought to be responsible for higher current source densities than the currents of action potentials. Others think of LFP as an artifact of bursts of action potentials^4, 5^. However, their fall off at 1/r in the case of monopolar sources and 1/r^2^ in dipolar sources makes for difficulty in explaining long-range transmission^1, 6^. Additionally, theta and gamma waves are reported to synchronize the spiking activity by varying inductance at the dendritic level^7^. But the mechanisms between volumetric synchronization and cellular synchronization have not been clearly bridged. Thus, the field is uncertain whether the resulting synaptic potentials, coincident EPSP’s, are evoked solely by simultaneous spiking or spiking plus axonal theta oscillations. Certainly, EEG signals require volume conduction of the LFP, but do not rule out the possibility that subthreshold oscillations in axonal membrane potential coordinate the synchronized presynaptic transmitter release, causing an LFP; we propose that axons carry the oscillatory signal in addition to spiking. Because LFPs are typically understood as voltages from volume conduction, we use the term “voltage oscillations” or, more specifically in cases where appropriate, “theta oscillations” to refer to the electrophysiologic activity from single axons.

In this study, we focus on the source of voltage oscillations in the theta range of 4-10 Hz because of their association with learning and memory for episodic and place memory. The first association of theta EEG with fear memory was reported by^8^, although isolated theta EEG recordings in rodents were seen earlier. These associations led^9^ to stimulate at theta frequencies the axonal inputs to CA1, the Schaffer collateral and temporoammonic pathways in rat hippocampal slices to produce long-term potentiation (LTP) in the CA1. The stimulation with an extracellular electrode potentiates bundles of axons from the CA3 and entorhinal cortex to enhance the firing of a population spike in the pyramidal layer of CA1. This raised the question to us of whether these axons naturally carry theta oscillations.

Beyond animal studies, a review of human studies established that theta oscillations support associative episodic and place memory through synchrony between the entorhinal region of the medial temporal lobe and the hippocampus^10^. Herweg et al. concluded with an outstanding research question that we have begun to answer, “Are changes in local theta power and inter-regional connectivity or synchronicity inherently linked?”

We have approached the problem of axonal subthreshold voltages by channeling axonal growth through microfluidic channels or tunnels. Originally devised by Taylor et al., 3 μm high x 10 μm wide microfluidic channels fabricated in silicone rubber (PDMS) selectively grow CNS axons through these tunnels if the length of the channel was at least 450 μm long^11^. Many others have used this two compartment design including our lab in which neurons were isolated from two separate subregions of the rat hippocampus (DG and CA3) and shown to maintain their unique mRNA subregion specificity after 3 weeks in a common culture medium^12^. The microfluidic tunnels isolate individual axons in a high resistivity channel above extracellular electrodes and enhance the biological signal amplitudes^13, 14^. Placement of the PDMS microfluidic tunnels over a microelectrode array (MEA) permits monitoring the direction of propagation of action potentials travelling down the axon. Narrower channels of 5 μm were more selective for single axons^14^. Our first four-compartment device enabled reconstruction of the hippocampal circuit from entorhinal cortical (EC) inputs through to dentate gyrus granule neurons (DG) to CA3 to CA1 and completion of the loop back to EC^15^. Spike rates and burst rates between the four subregions were different, indicating independent information processing. In Lassers et al., 2023, we found that patterned stimulation sharpened the routing and made responses more reliable. In^16^, we added a fifth set of tunnels from the EC to the DG to model the perforant path. In response to theta-burst stimulation, we reported more axons fired similar spike patterns. The EC-CA3 pathway enabled faster spiking within bursts and more spikes within burst compared to the four-tunnel architecture^17^. We first looked in axons for spontaneous spindle oscillations (LFP) between these four subregions^18^ where we found strongest spindles in the CA3 axons that pass to the CA1 with timing independent of spiking.

Local field potentials (LFP) are altered in Alzheimer’s disease^19, 20, 21^, schizophrenia^22^, sleep^18^, depression^22^, cognitive learning^23^ and most clearly epilepsy^21^. However, whether these LFP’s are a cause or an effect of disease is unclear. An effect of disease is assumed because these 1-300 Hz oscillatory voltages are thought to arise from the coordinate summation of subthreshold presynaptic currents measured by EEG, and because of limits in spatial resolution. However, a cause of disease should be considered if certain functional elements of can be demonstrated (**Box 1**). Satisfying these criteria allows us to suggest that pervasive brain theta oscillations originate from axons that enable long-range inter-regional communication. These criteria provide evidence for spontaneous multiplexed axonal oscillations (LFP) between hippocampal subregions which together with spikes influence target spiking.

Box 1. Credible evidence for a functional role of axonal voltage oscillations

1) Sparsity: In contrast a volume conduction, strong axonal voltage oscillation will be present in a low percentage of axons
2) Specific source somata: In contrast to an origin in coordinated synaptic events, the voltage oscillations in an axon will be tightly synchronized to its source neuron at near zero latency (<0.05 ms) due to fast axonal conduction velocities.
3) Specific target somata: In contrast to volume conduction, we will find selective neuronal targets with mono-synaptic delays of a few ms from the presynaptic axon to the post-synaptic target neuron
4) Phase and amplitude dependent target neuron responses: In contrast to volume-conduction and coordinated synaptic events, target spiking or bursting will depend on the axonal theta phase and amplitude of the voltage oscillation. Feed-forward potentiation increases spike probability at strong synapses for inter-regional communication.

## METHODS

### Four Chamber, Five Connection Tunnel Device

In brief, a microfluidic device was molded in PDMS (**Fig. 1**;^16^). The device was comprised of four chambers for separate culture of subregions of the hippocampus. Each chamber was 30 mm^2^. The subregional chambers were connected by 3 x 5 x 400 μm (HxWxL) microfluidic tunnels. In each array, 67 tunnels connected each subregion with four of those tunnels monitored by electrodes, excluding the EC-CA3 cross connection that had 11 tunnels with five tunnels monitored. These dimensions favor mostly one, two and three axons per tunnel (Narula 2017). Resistance within the tunnels was measured as R=ρL/(W×H), where ρ is the resistivity of the Neurobasal medium, 90 Ωcm^13^. This yields a resistance for the tunnel of 24 MΩ, far higher than the subregional electrode resistance 0.2 MΩ^13^. Therefore, these axonal tunnels produce high amplitude signals, well isolated from neighboring tunnels and the bulk subregion from which they originate. The device was affixed to a Multichannel Systems MEA120 1100 (Multichannel Systems, Reutlingen, Germany) for amplified recordings of spontaneous activity on the 120-electrode microarray.

**Fig 1.**
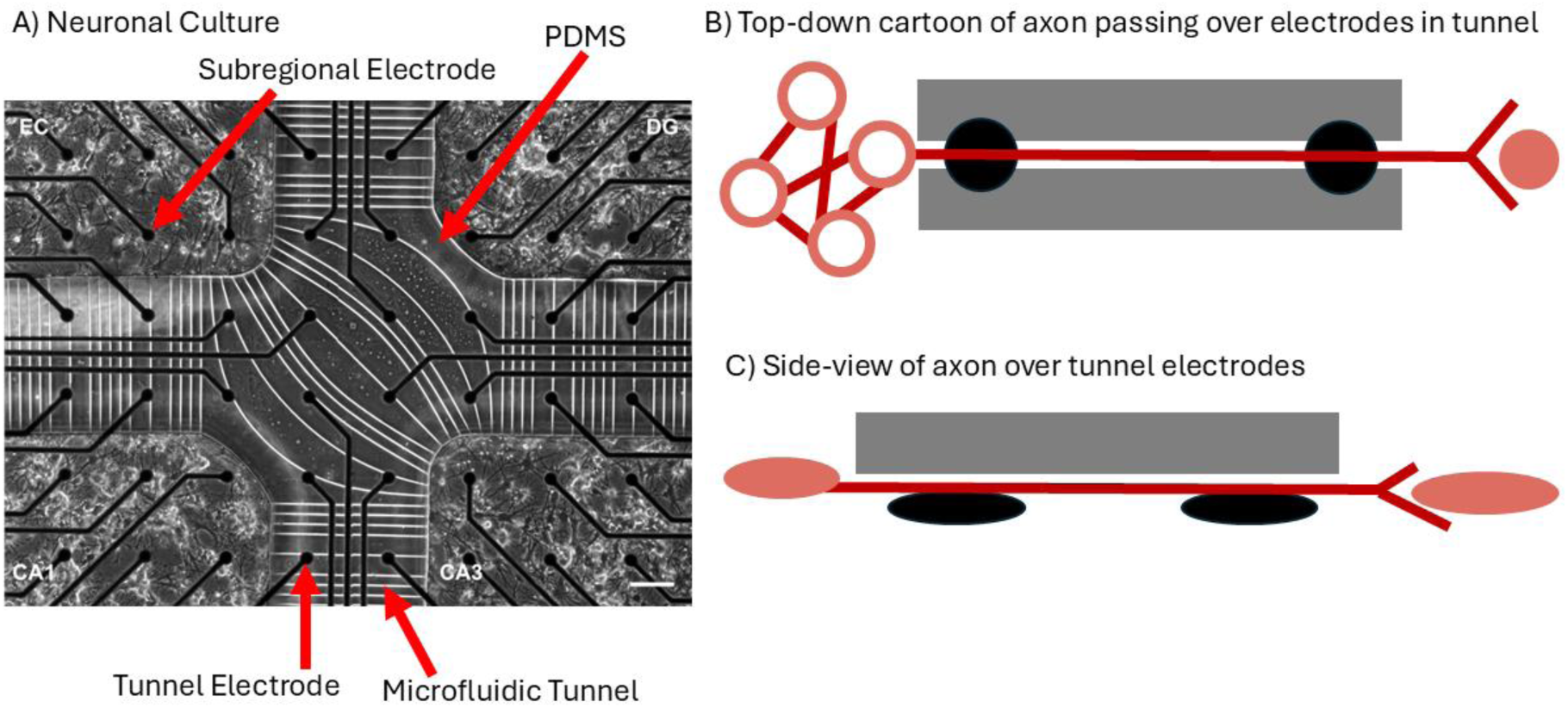
Five sets of tunnels enable axons to connect specific subregions of a hippocampal network. **A**) The four hippocampal subregions of entorhinal cortex (EC), DG, CA3, and CA1 were isolated and plated into device chambers. Axons grow through the micro-fluidic tunnels 3 μm high by 5 μm wide and 400 μm long (white lines) for the straight tunnels. The curved tunnels between EC and CA3 range from 560 to 980 μm long. **B**) Top-down diagram of an axon passing through a narrow high-resistance microfluidic tunnel over extracellular electrodes. **C**) Side view of an axon passing through a microfluidic tunnel, contacting electrodes.

### *In vitro* hippocampal neuronal network culture

Neurons were isolated and cultured from postnatal day 4 Sprague Dawley rat pups under anesthesia as approved by the UC Irvine Institutional Animal Care and Use Committee (IACUC)^16^. Briefly, entorhinal cortex (EC), dentate gyrus and hilus (DG), CA3, and CA1 including the subiculum were microdissected and cultured for 21 days in NbActiv4 medium^24^; Transnetyx BrainBits, Springfield, IL).

### Acquisition and processing of electrophysiological activity, spike sorting and direction of axonal information transfer

Spontaneous activity was analyzed from six networks (n=6 arrays) using MC_Rack (Multichannel Systems) and custom MATLAB software at a sampling rate of 25 kHz at 37 °C for 5 min. over the range of 0.5-50 kHz. Arrays with less than 80% active tunnels or with poor growth in one of the subregions were rejected for recording. Methods for spike sorting and determining the direction of axonal communication were described in our previous publications^15, 18, 25^. In brief, axonal spikes were sorted using Wave-Clus^26^ with sensitivity set to 5 S.D. of the signal for the tunnels and subregions^15^. Feedback and feed-forward direction of spike propagation was determined by the temporal delay in propagation of action potential over two electrodes spaced at 200 μm using our normalized matching index (NMI) algorithm^15^. Electrodes were considered for evaluation if the spike rate was at least 0.4 Hz averaged over the whole 300 second recording. We examined theta activity in axons via spectrograms generated by MATLAB’s continuous wavelet transform toolbox. After down sampling to 2500 Hz, the Morlet wavelet filter bank was set for the frequency ranges of 3-300 Hz at 10 voices per octave.

### Spike contribution to theta power

We wanted to evaluate whether theta oscillations were simply an artifact of repeated spiking. Spike cutouts from recorded signals were repeated at theta burst frequencies and zero-padded to 2 seconds for spectrographic analysis. We compared the power in the spectrogram for a single spike, three spikes pasted at theta timing of 100 ms and three trains of five spikes spaced at 200 Hz with trains repeated at 5 Hz. The spectrogram of each signal was computed for 3-300 Hz and summed over 4-10 Hz for total theta power. The summed magnitudes of the signal were normalized for frequency to reduce the over representation of higher frequency signals. The Hilbert transform was used on the theta filtered signal to visually represent the power of each signal.

### Prevalence of Axonal theta amplitudes

Theta amplitude was determined by filtering raw data collected from microelectrodes using a zero-phase 8^th^ order Butterworth filter for 4-10 Hz and down sampled to 1000 Hz. From the minimum to maximum theta amplitude of an interregional axon, a 100 log-binned histogram counted all Hilbert-transformed amplitudes in a single tunnel. Axons were considered high amplitude if they were above the interregional voltage mean plus one standard deviation for more than 10% of the recording (>30 sec). The integrated high amplitude axon voltages were averaged and compared by ANOVA with HSD correction for multiple sample testing.

### Identifying Source-Axon-Target Relationships

The goal was to determine the neuronal source of the axon recorded in the tunnel and the corresponding subregional target neuron. To find the source, we iteratively determined which, if any, of the 18 electrodes in the upstream subregion recorded neuronal theta activity that correlated with axonal theta by MATLAB XCORR with a 200 ms window (one theta cycle). Significance of the correlation was calculated using MATLAB CORRCOEF. To find the target neuron(s), we determined which axonal theta amplitudes or phase angles significantly modulated spikes in the target during axonal theta amplitudes above 5 μV. The influence of axonal theta on the target was computed as a modulation index^27^.

### Significance of Axonal Phase Angle and Amplitude with Subregional Spiking

Subregional spikes were counted within thresholded periods when axonal theta amplitudes were above background noise (5 μV) and longer than 200 ms (2x the shortest theta cycle). Intervals were connected if they were less than 3 cycles apart^18, 28^. To establish a relationship between the angle of axonal theta with subregional spiking, the Tort Modulation Index (MI) was used to determine if distributions of instantaneous axonal angles at subregional spike times derived from the Hilbert transform deviated significantly from a uniform distribution^27^. In short, a binned probability distribution of spikes over a range of angles was created and normalized for probability. The binning was determined via the Freedman-Diaconis rule (n=20) where the angle was on a linear scale from –180 degrees to 180 degrees. The Kullback-Leibler distance derived from the Shannon Entropy was divided by the log of the number of bins to get the Modulation Index. Nonparametric statistical testing was performed through shuffling high amplitude axonal angles 1000x via randomizing the imaginary component of the FFT against the subregional spikes in those same time intervals and a p-value was obtained. To establish a relationship between the amplitude of axonal theta with subregional spiking, a log-log binned probability distribution of subregional spikes over a range of axonal amplitudes was created. The binning was set equal to the number of angle bins (n=20). To obtain the significance of the amplitude measurements, a log-log linear regression was fit over this distribution for an R^2^, slope, and p-value of the slope significantly different from zero. The optimal range for the regression in each case was found via a grid search for the highest R^2^ value with the start of the regression anchored at the lowest amplitude (5 μV).

### Mutual information between axonal voltage oscillations

For specificity among axons, we determined whether multiple axons carried the same theta signals. Mutual information was used to determine how much information was shared in axons communicating in the same direction^29^. Binning was determined by the Freedman-Diaconis rule (n=20). The amplitudes of the axonal Hilbert envelopes of theta waves as the basis for comparison. The range spanned from the minimum amplitude of an axon to the maximum amplitude and was logarithmically spaced. This ensured that coupling of an axon to an electrode did not affect its ability to be compared to another axon, in effect normalizing the amplitudes between two axons. Distributions of binned amplitudes were compared to see how much information was in the same bin between every axon pair in a subregion, computed in bits.

### General Linear Models for predicting subregional spiking

General linear models (GLM) were used to predict if the amplitude and phase features of theta in axons had an effect on subregional target neuron spiking. The MATLAB R2024 fitglm function was used to calculate a GLM for predicting subregional spiking during intervals of high amplitude theta (selected using thresholding method explained above). The GLM was computed on signals down sampled to 2500 Hz. The response variable was target neuron spiking. Spiking was coded as indices of logical 1’s. The logit link function was used to determine if the independent variables of theta amplitude and phase predicted target neuron spiking. Theta phase was transformed to the cosine of theta during high theta episodes. Cosine was chosen because it linearized the cyclical relationship of angle and had a stronger prediction of spiking than either angle in degrees or the sine of the angle alone. All other indices below the threshold were not considered. Additionally, we computed GLMs for bursting activity in target neurons. All indices in the burst bounds were defined as “bursting” vs “non-bursting” in the indices outside the burst bounds. We defined the GLM using the formula in Wilkinson Notation:

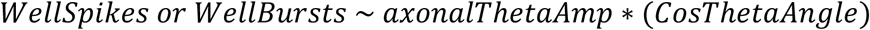

ThetaAmp was the z-scored amplitude of theta at each index above threshold. We defined bursts as a minimum of four spikes with a maximum of 50 ms between spikes.

### Statistics

Significance was determined as p<0.05. Data in bar graphs are means +/− S.E. with n independent observations. Bar averages were compared by ANOVA (unless this is specified above). Modulation indices (MIs) were subjected to non-parametric testing to determine significance. For amplitudes, indices within large amplitude theta regions were shuffled 1000 times and new MIs were calculated based on the shuffling. For angle, the phases were randomized using the custom MATLAB program phaseran^30^. The p-value was computed by summing the number of MIs that were randomly above the initial MI divided by the number of random permutations. A false discovery rate routine was also applied to the modulation indices and linear regressions to test the chance null-hypothesis.

## RESULTS

### A four-chambered device for reconstitution of the hippocampus with connecting axons

A custom PDMS device was used to accommodate separated subregions of the hippocampal formation into four chambers that contained cultured entorhinal cortex, dentate gyrus, CA3, and CA1 neurons from neonatal mice (**Fig. 1**). Five microfluidic tunnel systems connected these chambers including diagonal-cross tunnels which allowed for the formation of the perforant pathway from the EC to both the DG and CA3. Other tunnels allowed for connection of DG to CA3, CA3 to CA1 and CA1 to EC to complete a loop. Two electrodes spanning each tunnel allowed for the direction of information transmission to be determined based on spike times^17, 18, 25^. For reference, the alphanumeric code for each electrode is provided in **Supplementary Fig. 1**.

### Single axons transmit a rich repertoire of oscillatory voltage signals

**Supplementary Fig. 1.**
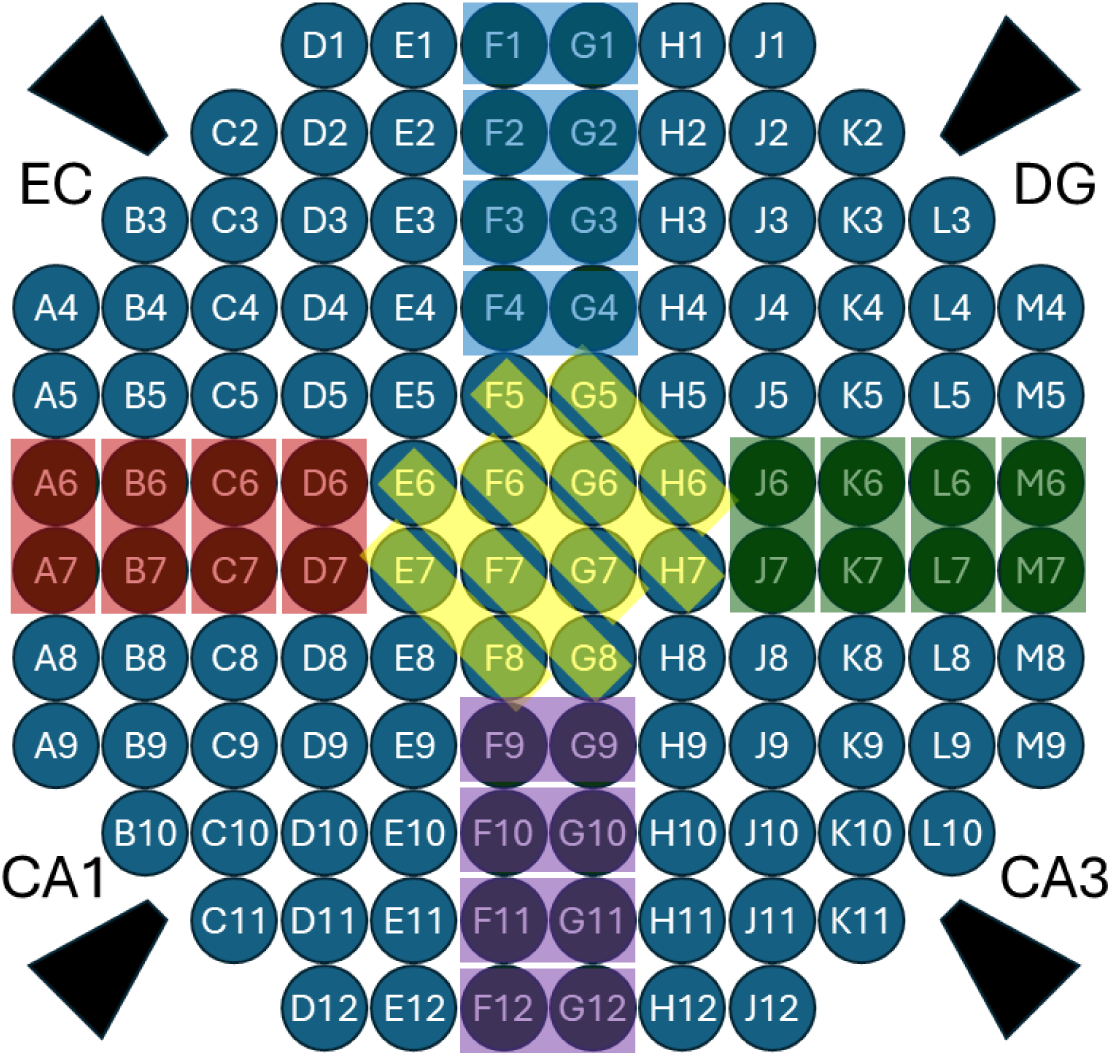
Electrode map with axon tunnels highlighted. We recorded from all 120 electrodes simultaneously so that activity in neurons in one subregion can be followed into some of the axons and through to some of their targets in another subregion. Grounding electrodes are marked by black parallelograms.

We recorded raw spontaneous axonal activity with coincident spiking and voltage oscillations at theta frequencies on an electrode with a contacting axon passing through each microfluidic tunnel. We examined frequency ranges from 3-300 Hz with 4-10 Hz theta, 10-16 Hz as spindle frequencies^18^, 30-100 Hz as low gamma, and 100-300 Hz as high gamma. Spikes were detected via filtering at 300-3000 Hz. **Fig. 2A** shows an axon from CA3 into CA1 (hereafter, CA3-CA1) with strong theta power, embedded spindle, low gamma frequencies, and weaker high gamma power. **Fig. 2B** shows the same signal filtered for theta waves and the overlayed Hilbert transform of the envelopes. The spike filtered signal in the same axon is shown in **Fig. 2C**, coincident with the theta oscillations. This leads to the question: are axonal voltage oscillations just artifacts of their spiking activity?

**Fig. 2.**
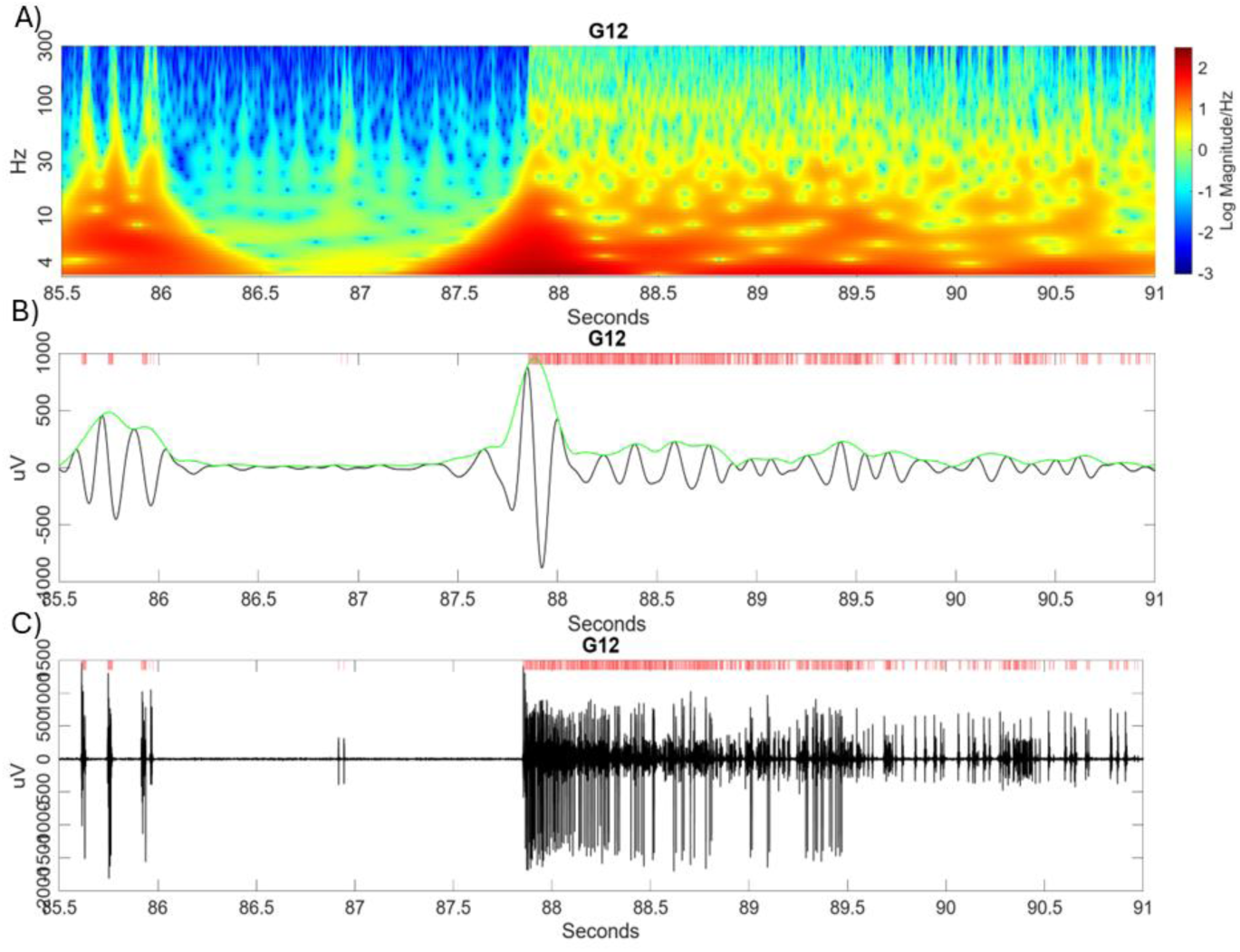
Rich repertoire of spontaneous single axon oscillatory signals with peak theta magnitudes corresponding to spike times. **A**) Frequency-magnitude spectrogram from 3-300 Hz for a single axon passing from CA3 to CA1 with strongest signals in theta (4-10 Hz) and low gamma (30-50 Hz). **B**) Raw signal from the same axon filtered for theta. Red ticks indicate timing of spikes in C). **C**) Same axon filtered from 300-3000 Hz to show a burst of spikes of action potentials corresponding to the start of the large amplitude theta wave.

### Axonal spikes at theta frequency contribute miniscule theta power

In order to determine how much theta power was attributed to the spikes, we compared a single burst of raw data (**Fig. 3A**), the corresponding spectrogram (**Fig. 3B**) and its theta filtered data (**Fig. 3 C**) to two different models of theta burst spiking. The first model in **Fig. 3D, E, F** is a paste-up of three spike cutouts separated by theta timing of 5 Hz (200 ms). The second model simulates a theta burst by separating 5 spike cutouts at 200 Hz (5 ms) as three bursts of spikes separated by 5 Hz (**Fig. 3 G, H, I**). In the 4-10 Hz range, the power of the spectrograph was summed to estimate the total power of the theta band frequency. The maximum amplitude derived from the Hilbert transform shows a 100-1000-fold decrease in signal amplitude (**Fig. 3 C, F, I;** note changes in Y axis). The summed power of the spectrograms from 4 Hz-10 Hz (**Fig. 3 J**) showed that the single spike model (not shown) and 5 Hz model (**Fig. 3 D, E, F**) contained approximately 1000x less power over the same time scale than the raw signal (**Fig. 3 A, B, C**). In the last bar in **Fig. 3 J**, the 200 Hz burst model summed from 4-10 Hz over the same time scale (**Fig. 3 G, H, I**) was 100x less powerful than the raw signal. We conclude that the 3 ms spike cutout contains virtually no theta power and that theta LFP activity exists together with spiking activity as another form of information.

**Fig. 3.**
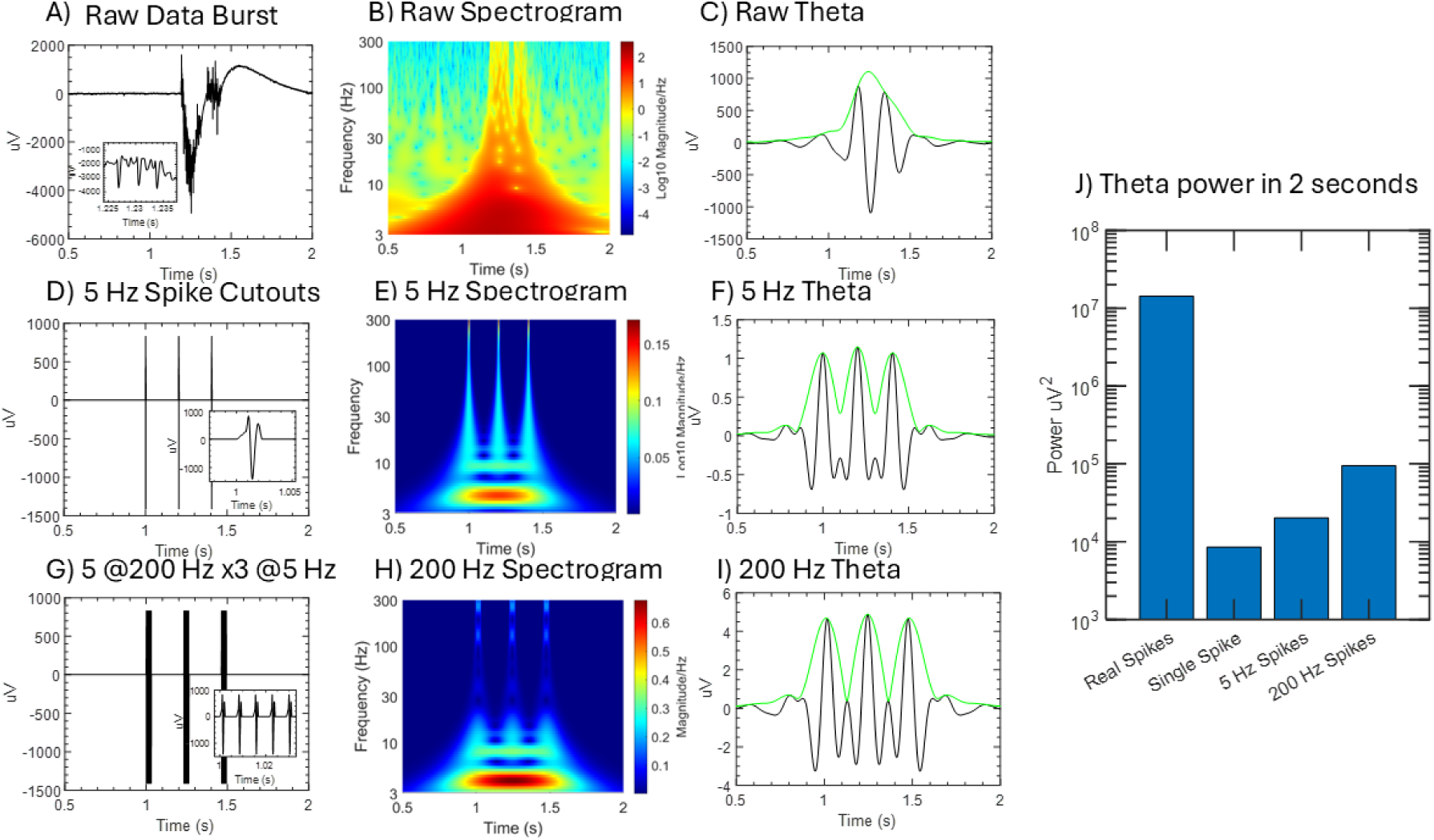
Spikes alone account for less than one percent of the theta power present in axons. **A**) A burst of spikes in raw data in an axon (FID 6 G12), **B**) the associated spectrogram from 3-300 Hz, and **C**) theta filtered data (black) with an overlayed Hilbert transform of the envelope (green). The power was calculated by thresholding the data above 1 SD of the Hilbert envelope and integrating the spectrogram from 4-10 Hz across those indices. **D**) Three ms of a single spike was cutout from the spike filtered signal in A (300-3000 Hz) and was repeated three times at 5 Hz theta spacing. **E**) The associated spectrogram (note lower maximum color scale) and **F**) theta filtered signal (note lower maximum y-axis). **G**) A burst of five cutout spikes were spaced at 200 Hz with three repeats spaced at 5 Hz. **H**) The corresponding spectrogram and **I**) the theta filtered signal of this theta burst model. **J**) The log-scaled power of theta for each model compared to the raw data (single spike not shown).

### Strong theta is sparse and strongest in CA3-CA1 axons

To transmit specific information, high power theta waves in axons should be sparse. We examined the log-normal distribution of theta power for every axon with spikes (**Fig. 4**). High axonal power was classified as any axon that had signal 1 SD above the mean (about 10 µV in all subregions) and above that threshold for at least 10% of the recording time (10% of 300 s = 30 s total). Twenty-two percent of axons had strong theta in communicating axons (**Fig. 4 A-E**). CA3-CA1 axons had the highest theta amplitudes with 3/14, 21% axons having strong theta above threshold. This thresholding method was applied to other axons which resulted in axons classified as high amplitude within EC-DG 3 of 12, DG-CA3 4 of 15, CA1-EC 2 of 12 and EC-CA3: 2 of 11. The highest power CA3-CA1 axons were significantly different from EC-DG, CA1-EC, and EC-CA3 (**Fig. 4 F**). These results suggest specificity in the sparse architecture that emerges in high-power communication between subregions.

**Fig. 4.**
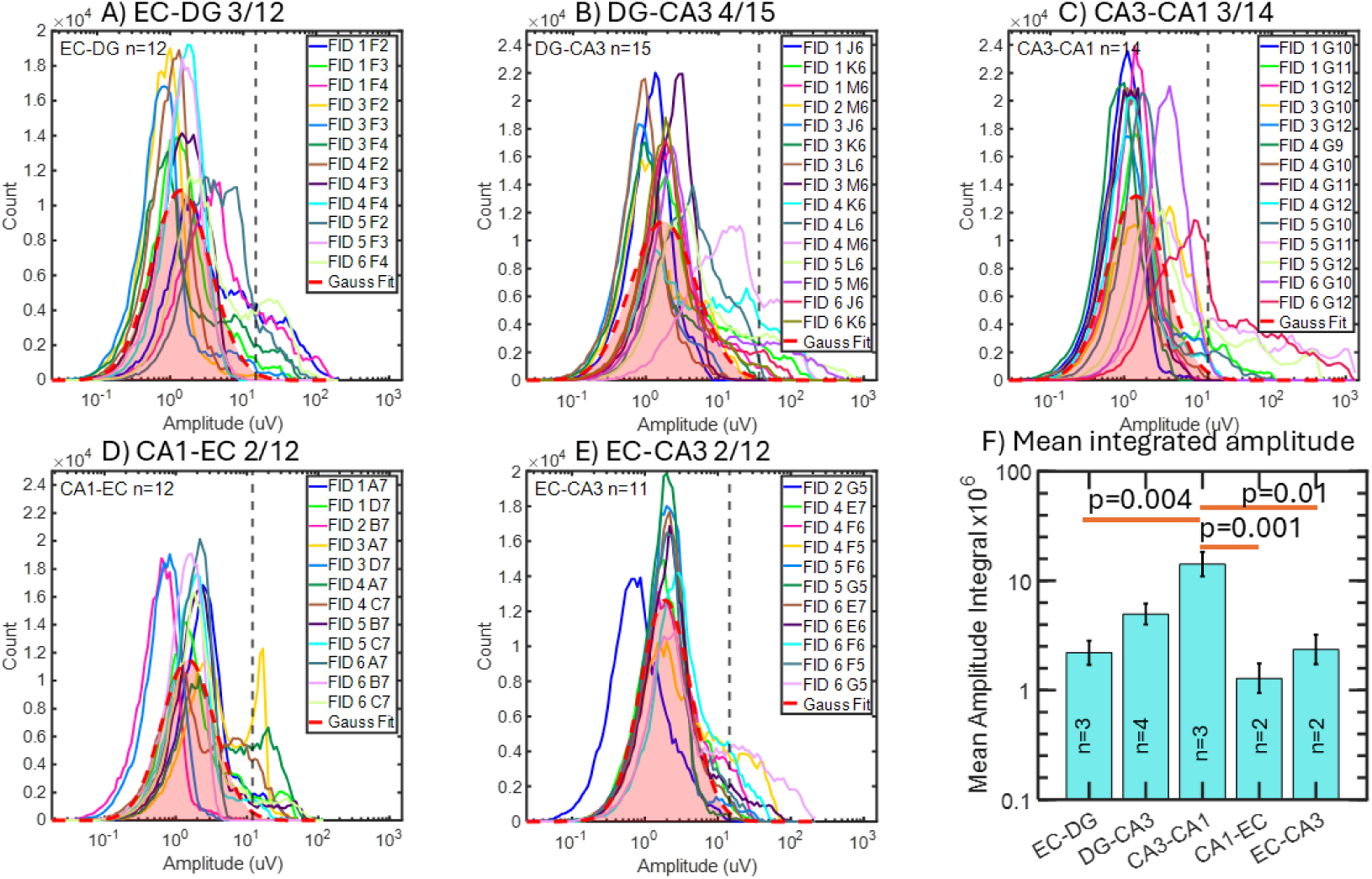
Among all axons traversing between subregions, the highest theta amplitude waves were from a few sparse CA3-CA1 axons. The Gaussian fits of all amplitudes of all axons in a subregion are indicated as a dashed red line. Axons were demarcated as high amplitude when their amplitudes crossed a threshold of mean+1 SD for at least 30 seconds (10% length of signal). High amplitude theta activity in the **A)** EC-DG, **B)** DG-CA3, **C)** CA3-CA1, **D)** CA1-EC and **E)** EC-CA3 was between 17%-26% of all axons (displayed in subtitles as number of high amplitude/all axons in all arrays). **F)** The relative differences in amplitude between the high amplitude axons integral. Only CA3-CA1 axons had significantly higher amplitudes.

### Axonal theta phase and amplitude evoke spiking in subregional target neurons via a power-law relationship

To show further specificity, a single source neuron should be associated with each axon with theta waves and should affect the spiking of a limited set of target neurons, due to branching axons. Since we only have 18 electrodes that monitor the source or target well and several hundred neurons in each well, there’s a chance we will not detect the direct source or target neuron or that we will only be able to see a relationship with a grandparent source or daughter target. Starting with a CA3-CA1 axon with high gamma power, we searched every neuron in CA3 for one whose theta oscillations coincided with those of the axon. **Fig. 5A and B** show such a relationship that is not perfect, suggesting a CA3 grandparent neuron connects to another neuron whose axon we recorded. Further, spikes in a possible CA1 target neuron (**Fig. 5C**) were closely aligned with the high amplitudes of theta oscillations in the axon. We focused on feed-forward axons to examine their effects on one or more of the possible 18 target neurons that we could monitor. Due to the potential that feedback axons inhibit targets, we cover only feed-forward axons in this paper; feedback inhibition would require more complex algorithms to categorize. By cross correlation of the theta oscillations in the putative source neuron with an emanating axon, **Fig. 5D** shows a strong peak with a delay of 21 ms. This long delay suggests that the source neuron was a grandparent that connects through at least two synapses before reaching the primary source of the axon that was monitored.

**Fig. 5.**
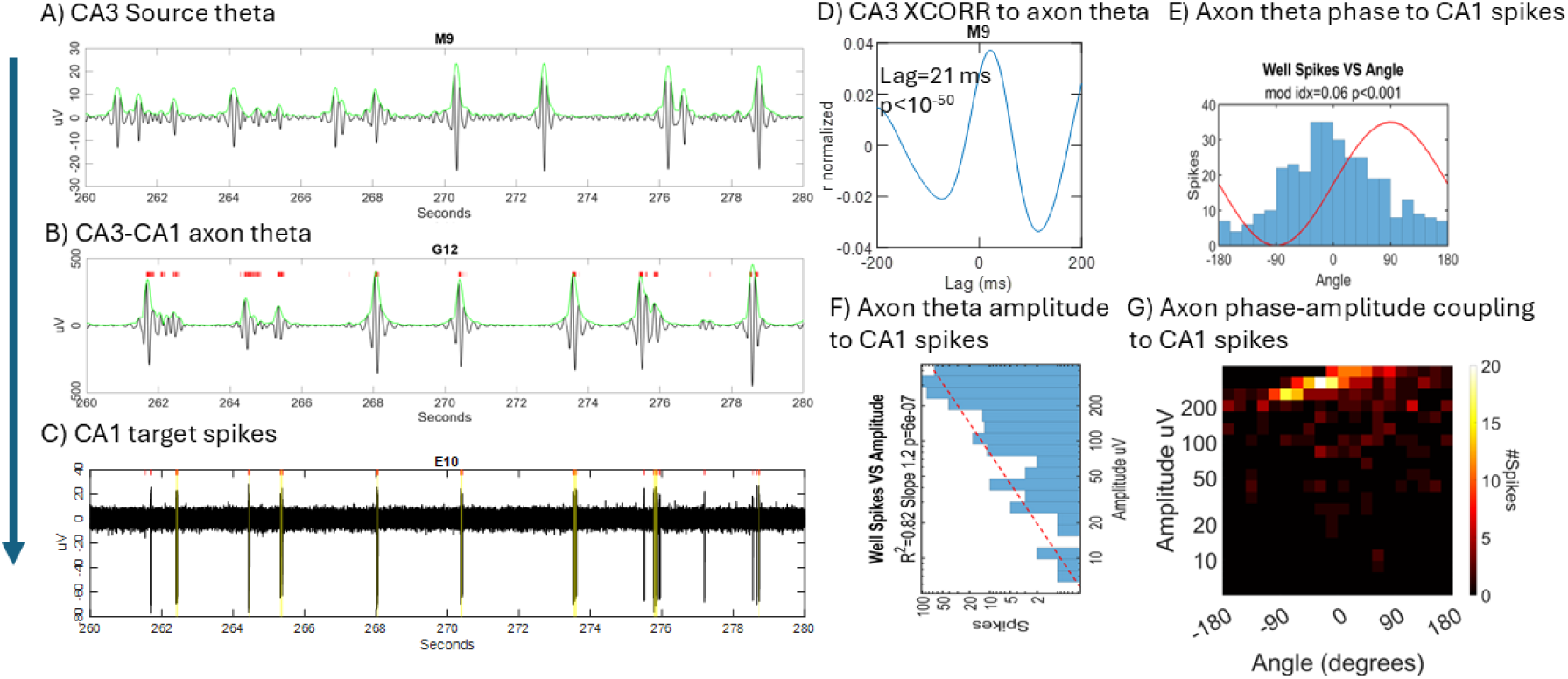
Theta CA3 source neuron to CA3-CA1 feed-forward axon evoked synchronized target neuron spiking. **A**) A single source neuron in CA3 identified by correlation with theta in axon. Theta signal in black, Hilbert envelope in green. **B**) Axonal theta oscillations and filtered spikes (red ticks) on G12. **C**) Target neuron in CA1 identified by correlation of spikes (red ticks at 5 SD) with timing of theta in axon. **D**) XCORR to demonstrate theta correlation in CA3 source neuron from electrode M9 to axon G12. M9 likely a grandparent since lag was large and correlation not perfect. **E**) Correlation of G12 axonal theta phase angle with spiking in E10 target neuron. Peak near zero at upswing from negative to positive amplitude. Red sine wave plotted to illustrate phase. **F**) Correlation of axonal theta amplitude histogram with spiking in CA1 E10 target neuron. Peak at 355 μV. Red linear regression and correlation coefficient noted. **G**) Phase-amplitude coupling of axonal theta to CA1 target neuron spiking.

From the distribution of evoked spikes in the subregion that corresponded to instantaneous amplitude and angle measurements from axonal theta (**Fig. 5C**), we calculated the log-log linear regression for amplitude (**Fig. 5F**), and the modulation index of theta wave angles (**Fig. 5E**). The modulation index was used to compare the actual distribution of spikes with phase angle to a shuffled distribution of angles and derive a non-parametric statistic. It is important to note that the amplitude has a log-log relationship with evoked spiking in the CA1 subregion as this follows established power-law relationships found in other neural systems. **Fig. 5G** shows the relationship between angle and amplitude for subregional spiking. Spiking in the subregion was modulated by the maximum dV/dt when theta angle crosses 0 degrees and derivative of the change in theta amplitude was the highest. This example and others not shown indicate specific functional effects on target spiking at times of high amplitude and specific theta phase in single axons. As a negative control, **Supplementary Fig. 2** shows a control case when there was poor correlation between axonal theta phase and subregional spiking, but theta amplitude was still prominent. **Supplementary Fig. 3** shows a control case when there was poor correlation between axonal theta phase, and amplitude with subregional spiking.

**Supplementary Fig. 2.**
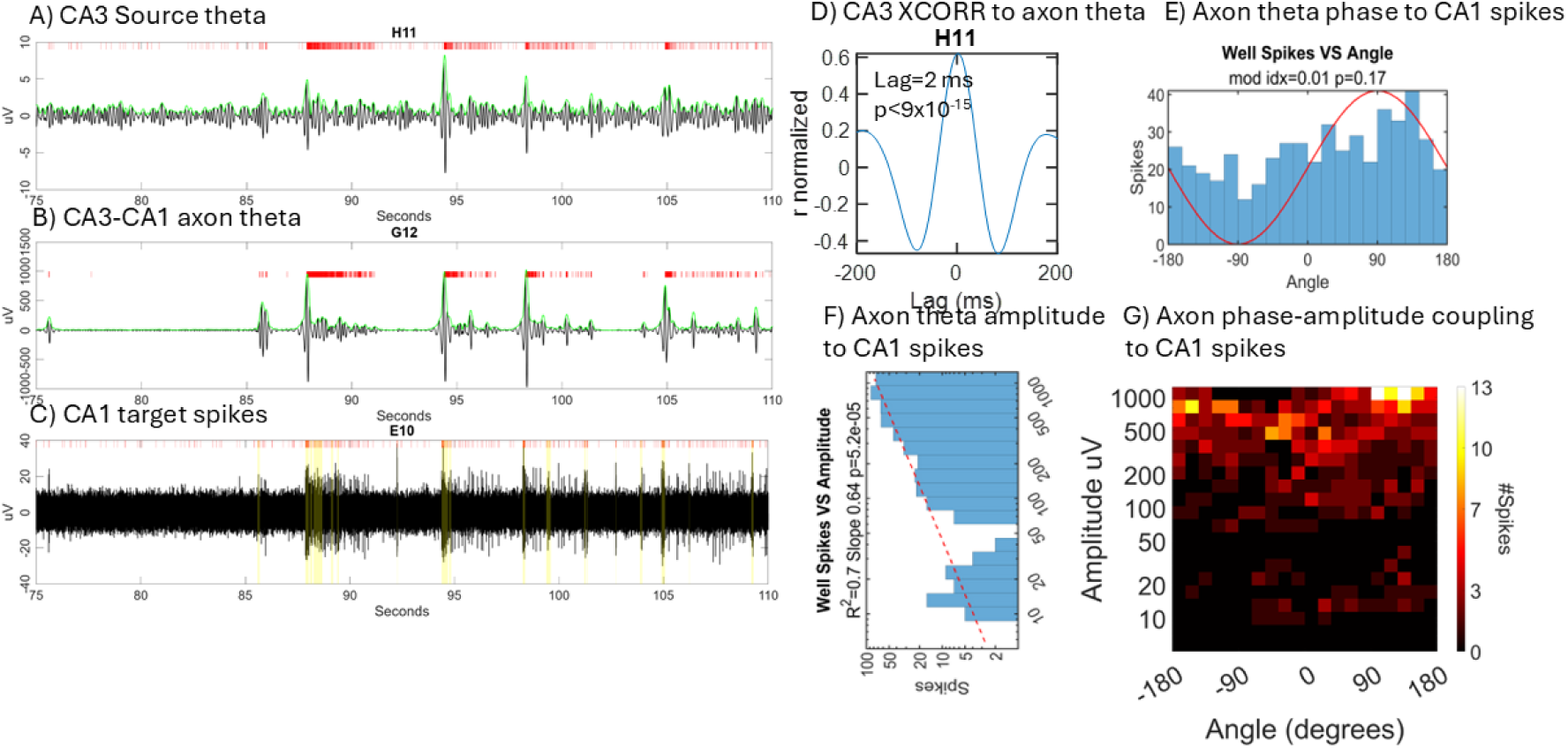
Negative control of CA3-CA1 axon that failed to evoke spikes at specific theta phases but was still amplitude coupled to spikes in target CA1. **A**) A single source neuron in CA3 identified by correlation with theta in axon. Theta signal in black, Hilbert envelope in green. **B**) Axonal theta oscillations and filtered spikes (red ticks) on G12. **C**) Target neuron in CA1 identified by correlation of spikes (red ticks at 5 SD) with timing of theta in axon. **D**) XCORR to demonstrate theta correlation in CA3 source neuron from electrode H11 to axon G12. **E**) No correlation of G12 axonal theta phase angle with spiking in E10 target neuron. **F**) No correlation of axonal theta amplitude histogram with spiking in CA1 E10 target neuron. **G**) Poor phase-amplitude coupling of axonal theta to CA1 target neuron spiking.

**Supplementary Fig. 3.**
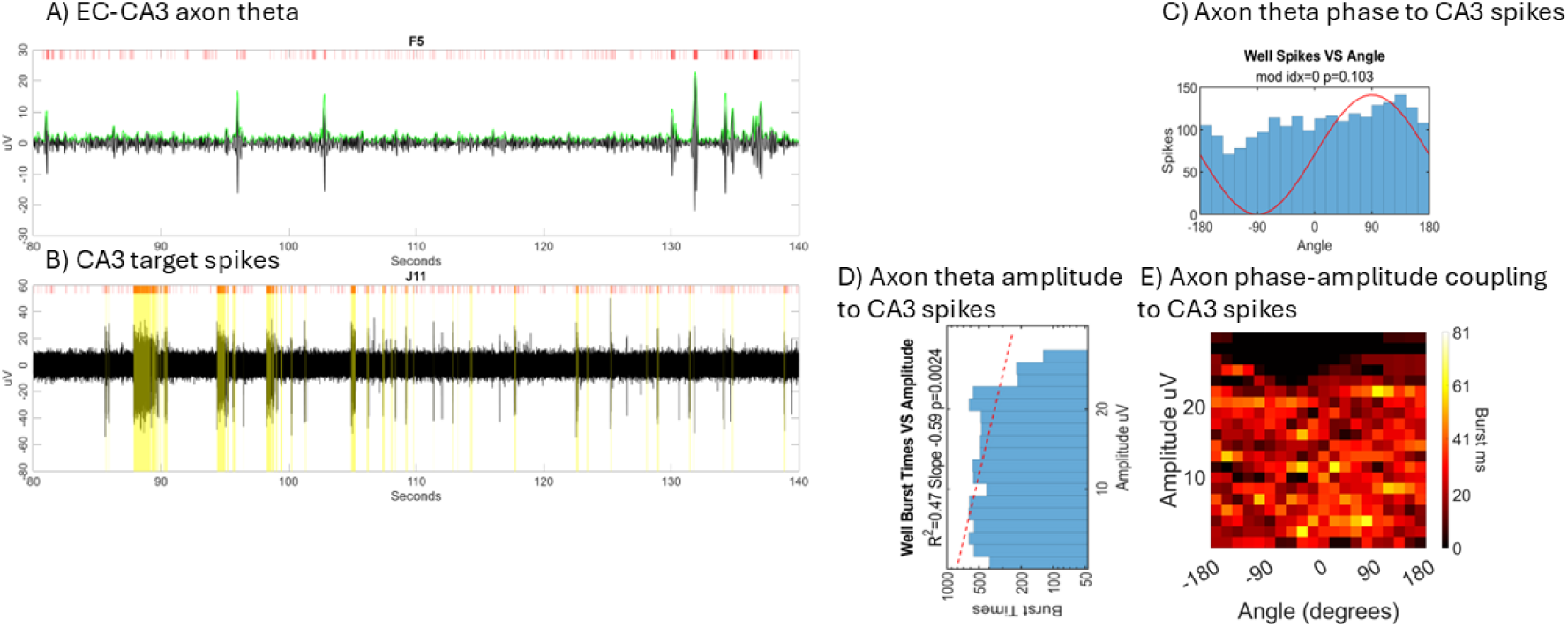
Negative control of neither theta amplitude nor angle in an EC-CA3 axon affecting target spikes. **A**) Axonal theta oscillations and filtered spikes (red ticks) on F5. **B**) Non-target neuron in CA3. **C**) No correlation of F5 axonal theta phase angle with spiking in J11 target neuron. **D**) Negative correlation of axonal theta and subregional spiking. **E**) Heatmap relating phase and amplitude.

To generalize the phase angle dependance of somal spiking for multiple axons by subregions and validate specificity, we predicted a peak angle would emerge for each subregion. **Fig. 6** shows three broad classes of peaks. **Fig. 6 A, B, C and D s**how all theta peaks at the upswing of the theta wave, near zero degrees for EC-DG axons, **DG-CA3, CA3-CA1, and CA1-EC** axons, respectively. Among the interregional axons with multiple targets, the peaks for CA3-CA1 were most prominent with a modulation index of 0.04. **Fig. 6E** for EC-CA3 axons shows a peak at the downswing to the theta wave at 135 degrees. Like the second peak in **Fig. 6C for CA3-CA1, Fig. 6F** shows that a single axon from EC-CA1 produced a sharp 63° maximum, near the peak of theta for evoked spikes in a single target; no other targets for EC-CA1 were found. Overall, our hypothesis of specificity is supported that theta angles specific for each subregion evoke spikes in their target neurons. This also suggests that different mechanisms specify these phase groups for evoking target neuron spiking. Next, we examine the probability of axonal phase-amplitude coupling to evoke spikes and bursts.

**Fig. 6.**
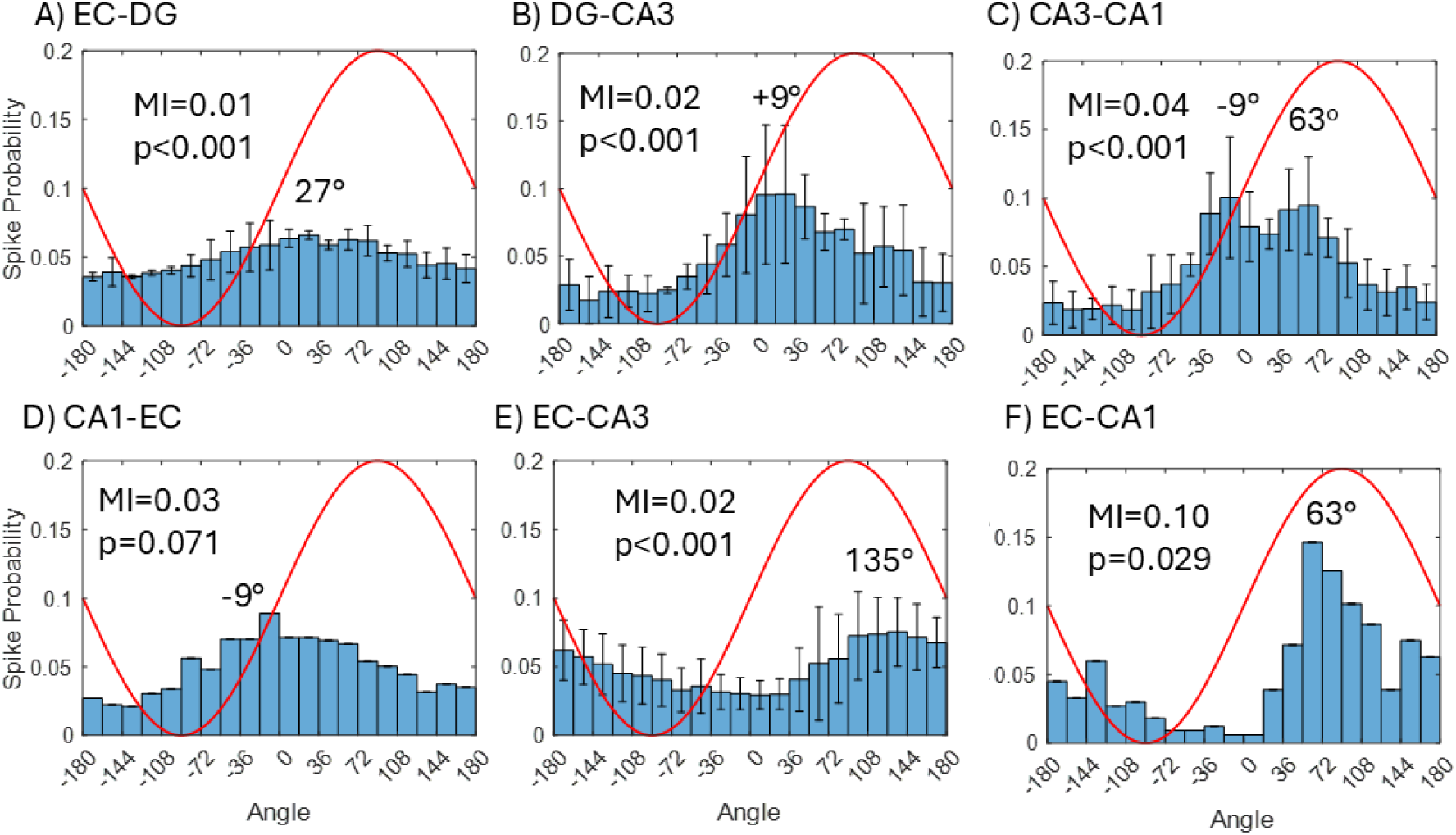
Two classes of theta angle in six feed-forward axon groups. The average axonal theta phase-angle evoking spiking was near zero for **A**) EC-DG peak at 27° (n=5), **B**) DG-CA3 peak at 9° (n=3), **C**) CA3-CA1 peak at both –9° and 63° (n=5), and **D**) CA1-EC peak at –9° (n=1). Peak spiking was evoked closer to 90° in **E**) EC-CA3 peak at 135° (n=23), and **F**) EC-CA1 peak at 63° (n=1), but only one axon-target pair could be found so error bars were not plotted.

### Interaction of axonal phase and amplitude for induction of target neuron spiking and bursting

GLM was used to quantify significant axon-target pairs of the axonal phase-amplitude coupling that affect subregional target spiking and bursting for multiple axons of each subregion type and multiple network arrays. The model was setup to predict subregional spiking based on the axonal theta amplitude, cosine of theta angle, and their interaction. Cosine of the theta angle was used to scale the angle from –1 to +1 which adds power in linearizing its effect for the GLM model better than Sine angle (**Supp Fig. 4**). We focused on the interaction of axonal cos (theta angle) and amplitude to determine their instantaneous coincident significance for association with spiking in each target neuron (**Fig. 7**). The significant interactions that contributed to spiking in each subregion: 9 of 20 (45%) possible targets for EC-DG feed-forward axonal connections (**Fig. 7 A**), 3 of 14 (21%) DG-CA3 connections (**Fig. 7 B**), 3 of 12 (25%) CA3-CA1 connections (**Fig. 7 C**), 0 of 11 CA1-EC connections (**Fig. 7 D**), and 2 of 34 (6%) EC-CA3 connections (**Fig. 7 E**). The individual contributions of amplitude (**Supp Fig. 5**) and angle (**Supp Fig. 6**) toward the prediction of spikes were evaluated. Both the axonal theta amplitude and the angle in the GLM alone rigorously supported the phase modulation index and log-log amplitude linear regression in which axons affect subregional spiking (**Fig. 5**).

**Fig. 7.**
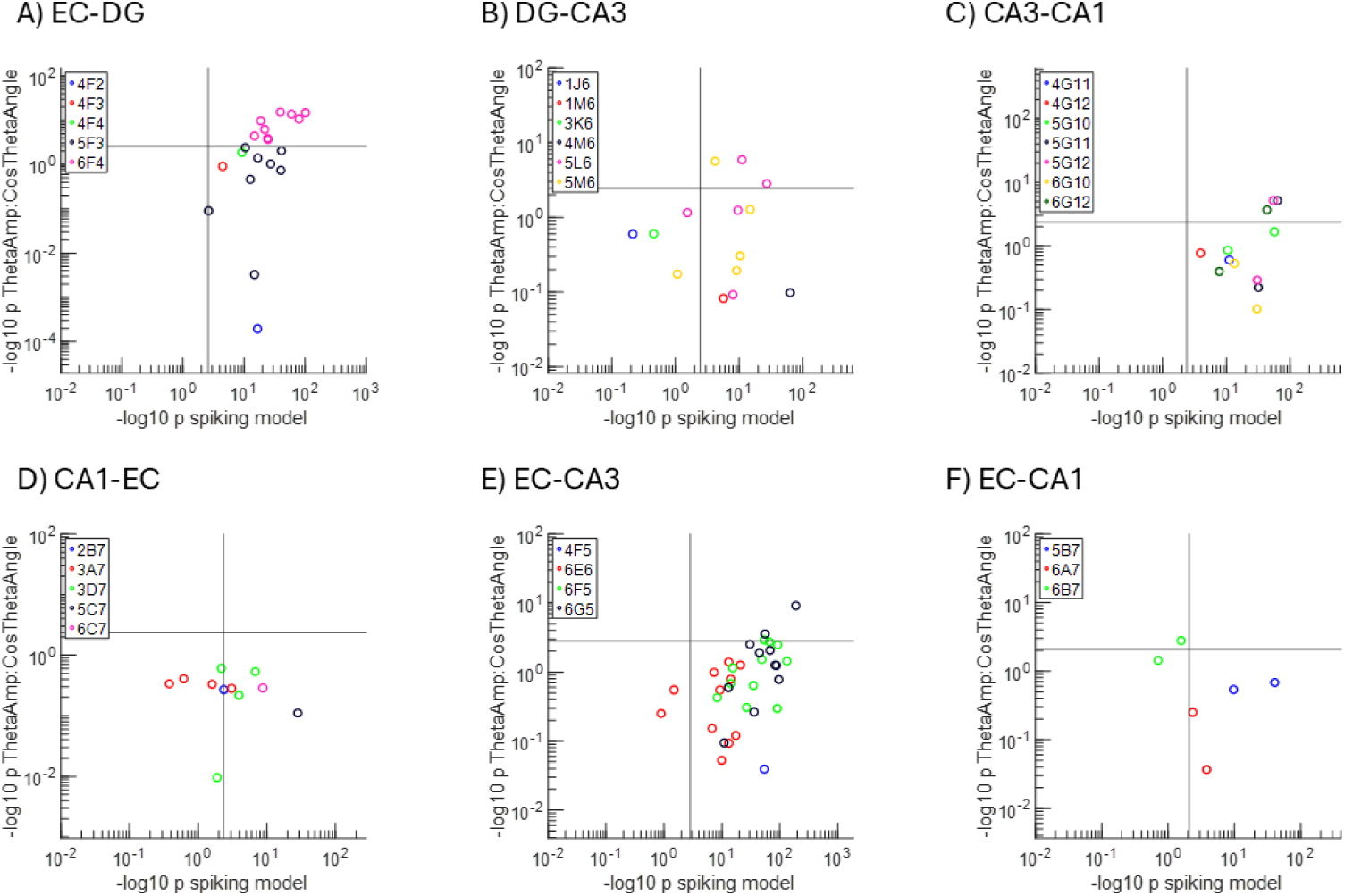
Influence of theta angle and amplitude interaction on the overall predictive power of subregional spiking shows sparse coding by theta axons. The frequency of significant axonal amplitude-angle interaction on subregional target neuron spiking over total spiking was in **A**) EC-DG 9/20, **B**) DG-CA3 3/14, **C**) CA3-CA1 3/12, **D**) CA1-EC 0/11, **E**) EC-CA3 3/34, and **F**) EC-CA1 0/6. Grey vertical and horizontal lines indicate the Bonferroni adjusted cutoffs for significance.

**Suppl. Fig. 4.**
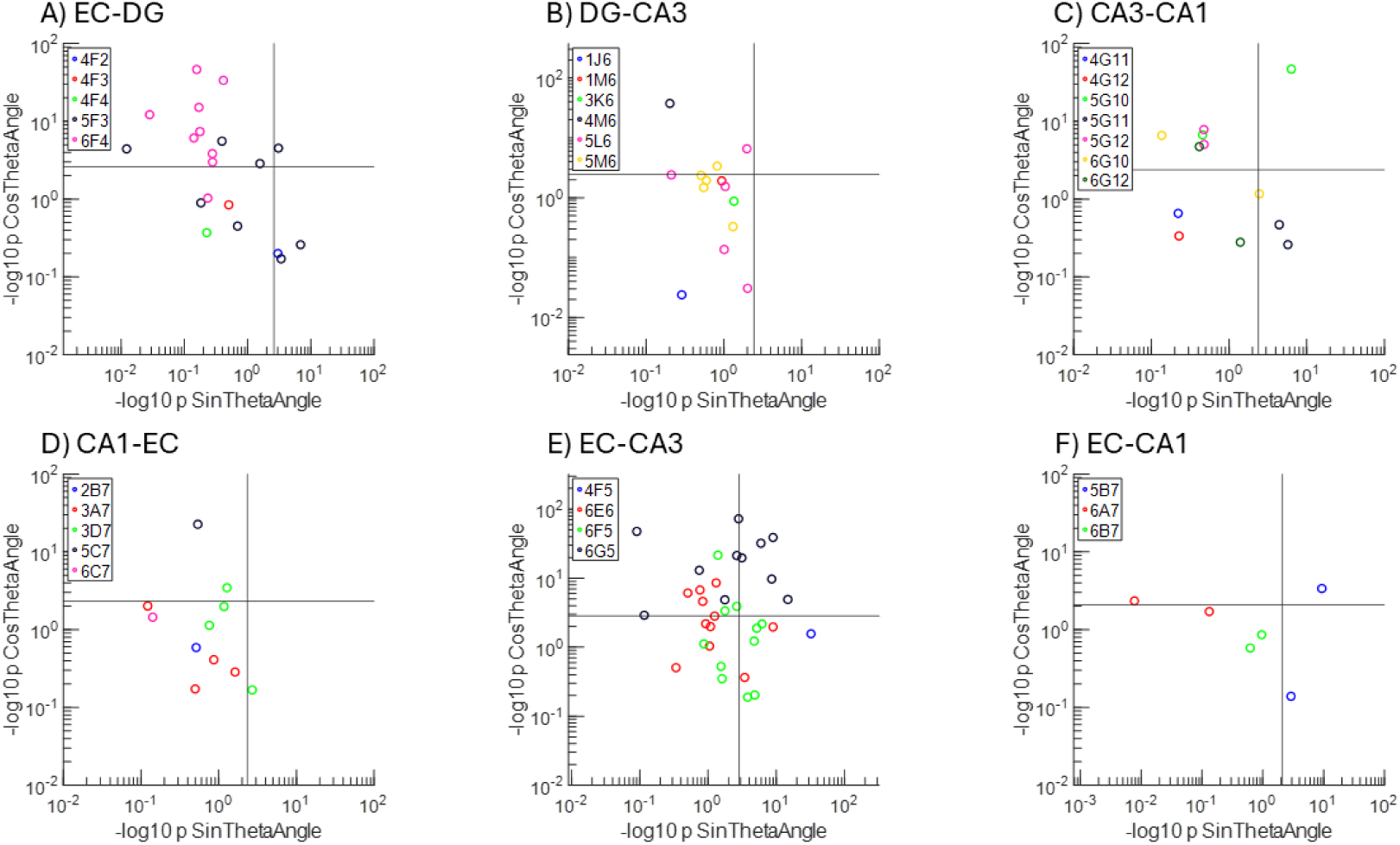
Cos angle predicts more target neuron spiking than Sin angle above and to the right of gray line significance cutoffs after Bonferroni multiple comparison adjustment. Five of six axonal inter-regions were more significant for cosine angles than sin angles as follows: **A**) EC-DG 12 cos vs 4 sin, **B**) DG-CA3 3 vs 0, **C**) CA3-CA1 6 vs 4, **D**) CA1-EC 2 vs 1, **E**) EC-CA3 19 vs 13, **F**) EC-CA1 2 vs 2. Each dot is the probability of spiking for one axon-target pair derived from the contribution of sine and cosine to the GLM.

**Supplementary Fig. 5.**
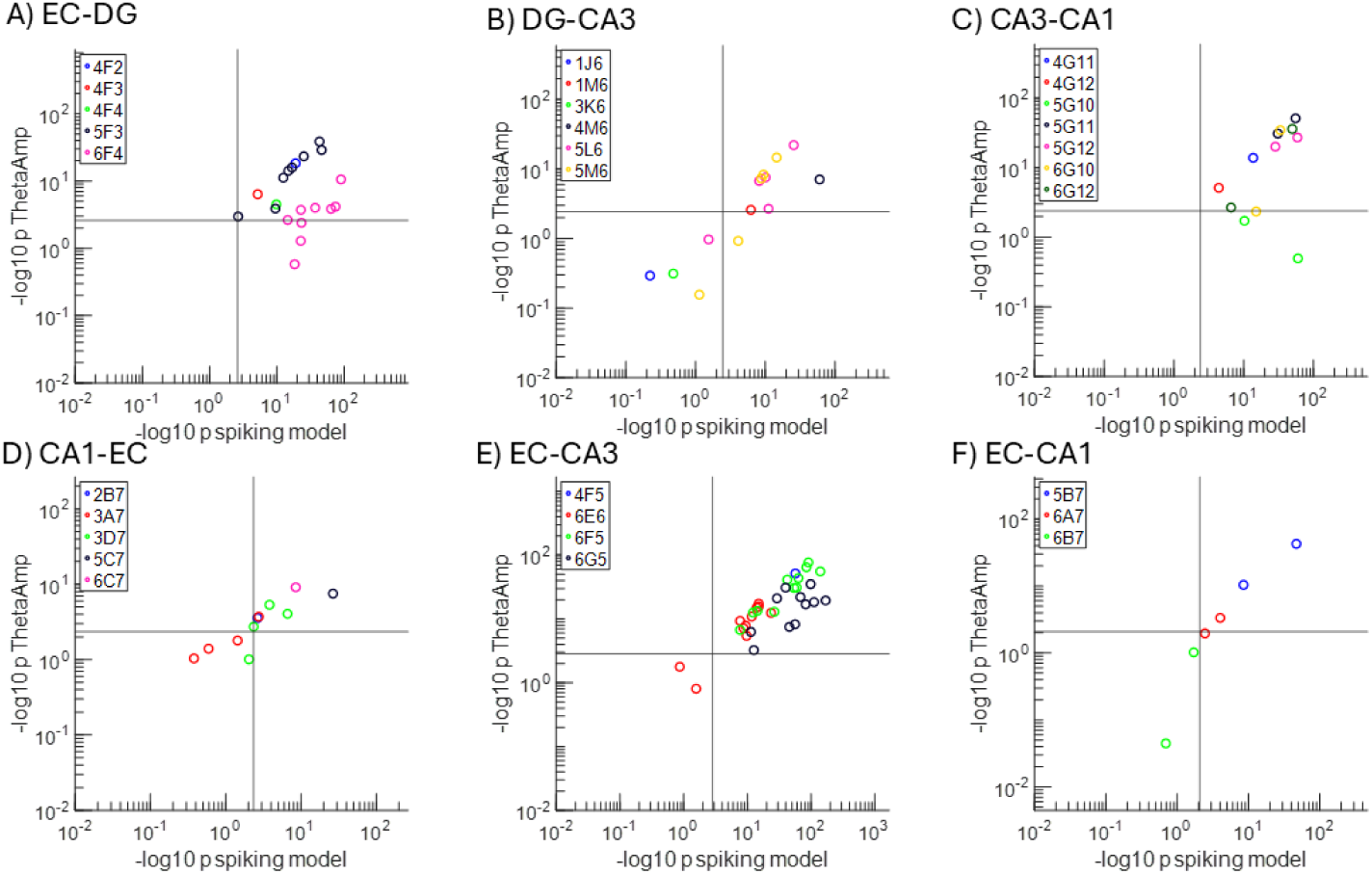
Amplitude alone was a strong predictor of subregional spiking with all inter-subregional axon amplitudes affecting spiking more than the interaction of amplitude and angle. **A**) EC-DG was significant for 17 of 20 axon-targets, **B**) DG-CA3 11 of 14, **C**) CA3-CA1 9 of 12, **D**) CA1-EC 8 of 11, **E**) EC-CA3 32 of 34, and **F**) EC-CA1 3 of 6. Grey vertical and horizontal lines indicate the Bonferroni adjusted cutoffs for significance.

**Supplementary Fig. 6.**
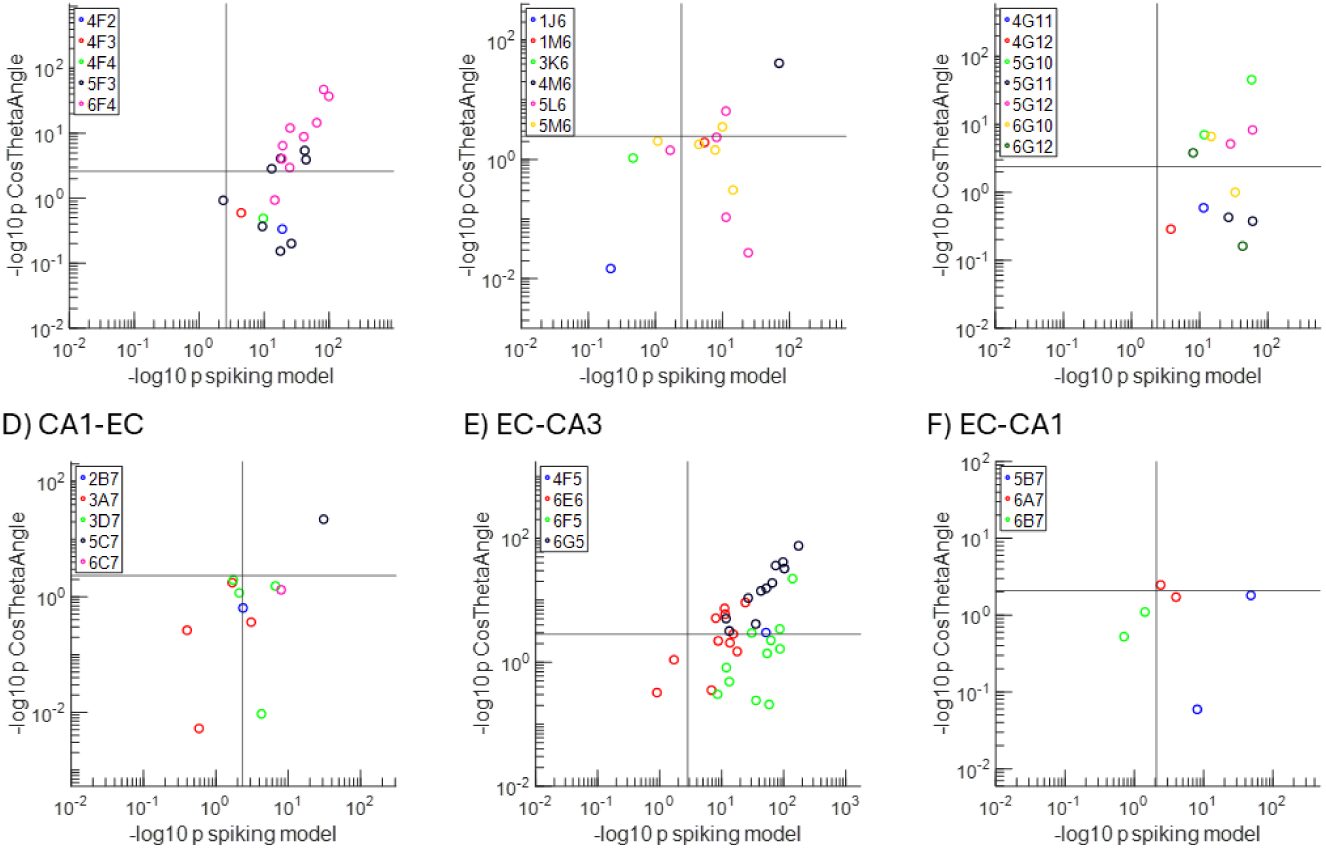
The contribution of cosine theta angle to the bursting prediction p-value. **A**) EC-DG was significant for 12 of 20 axon-targets, **B**) DG-CA3 3 of 14, **C**) CA3-CA1 6 of 12, **D**) CA1-EC 1 of 11, **E**) EC-CA3 20 of 34, and **F**) EC-CA1 1 of 6. Grey vertical and horizontal lines indicate the Bonferroni adjusted cutoffs for

### Theta waves evoke bursts more precisely than individual spikes

We next asked whether the GLM model indicated that axonal theta waves evoked bursts of spikes more reliably than individual spikes (**Fig. 8**). Compared to individual spikes, the interaction of axonal theta angle and amplitude was found to play a more significant role in predicting somal target bursting than spiking. The fraction of neurons responsive to the interaction of both axonal theta amplitude and the cosine of the theta angle that affected target bursting was 80% for EC-DG, 50% for DG-CA3, 83% for CA3-CA1, 64% for CA1-EC, and 73% for EC-CA3 (**Fig. 8 A-F**). **Fig. 8C2** shows a strong effect of theta phase and voltage amplitudes on the burst length in a target neuron. **Fig. 8G** shows that the interaction of amplitude with cosine of the theta angle predicts more target neuron bursts than neuron spikes in all subregions, ranging from 1.8 to 12-fold greater response fractions. The EC-DG and EC-CA3 axons have the largest number of targets, possibly due to the fan out nature of the EC subregion into these targets. For counts of individual target neurons affected by axonal amplitude and angle, **Supplementary Fig. 7** and **8** respectively show that axonal theta frequently evoked bursts of spikes.

**Fig. 8.**
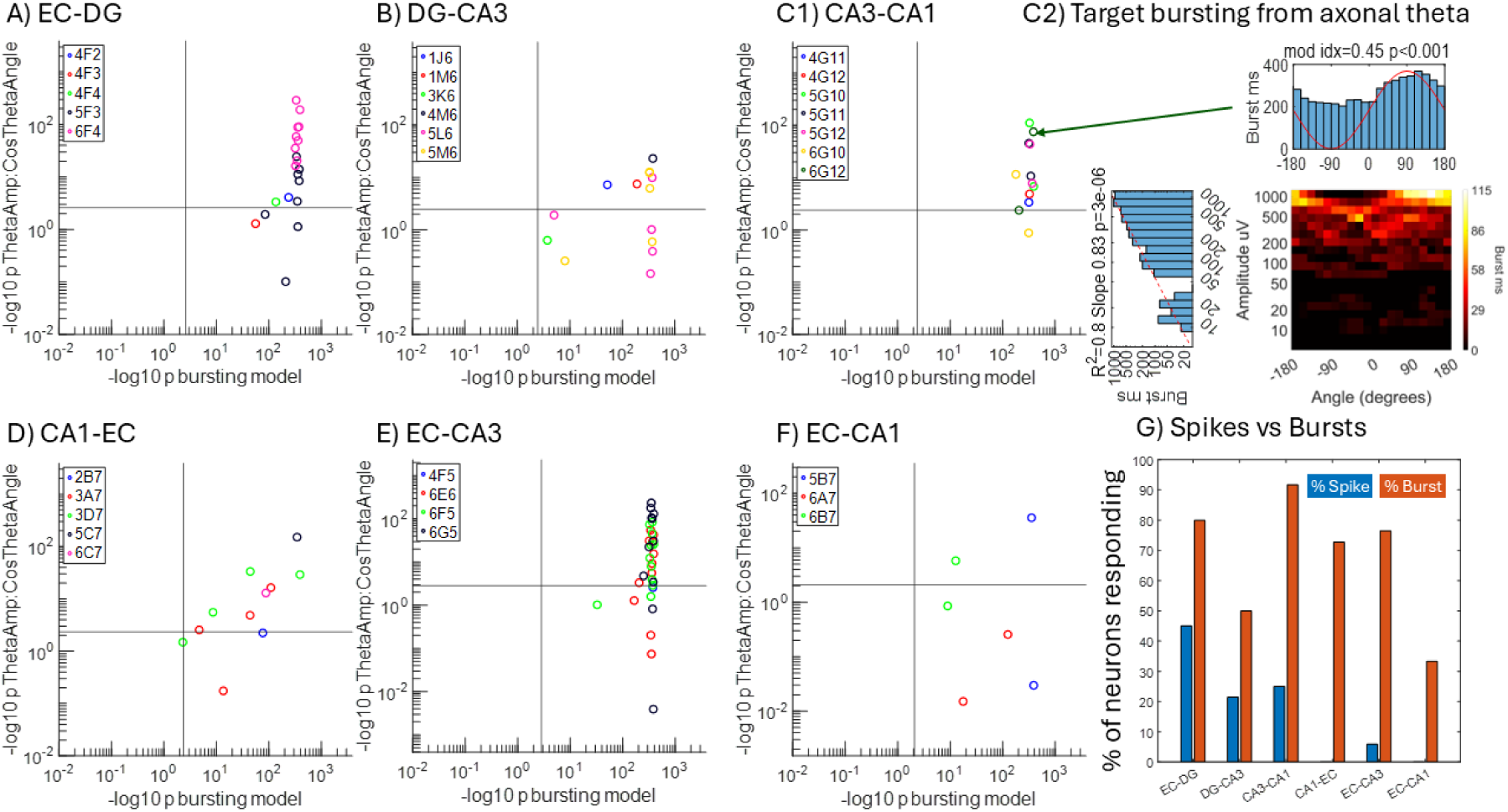
Subregional neuronal bursting evoked by axonal theta amplitude and angle interaction has a higher rate of correlation to subregional bursting than spiking. The subregional theta amplitude-angle interactions contributed to the overall prediction of bursting **A**) EC-DG 16 of 20 targets, **B**) DG-CA3 7 of 14, **C**) 11 of 12, **C2**) An example of a single CA3-CA1 axon theta phase-amplitude coupling for target bursting. Spiking increases linearly in log-log space with axonal amplitude and bursting was evoked after maximal dV/dt was reached during the rising phase of theta. **D**) CA1-EC 8 of 11, **E**) 26 of 34, and **F**) 3 of 6. G) Relative axon-target percentages with significant amplitude-angle interaction terms that predicted bursts was higher than those that predicted spikes.

**Suppl. Fig. 7.**
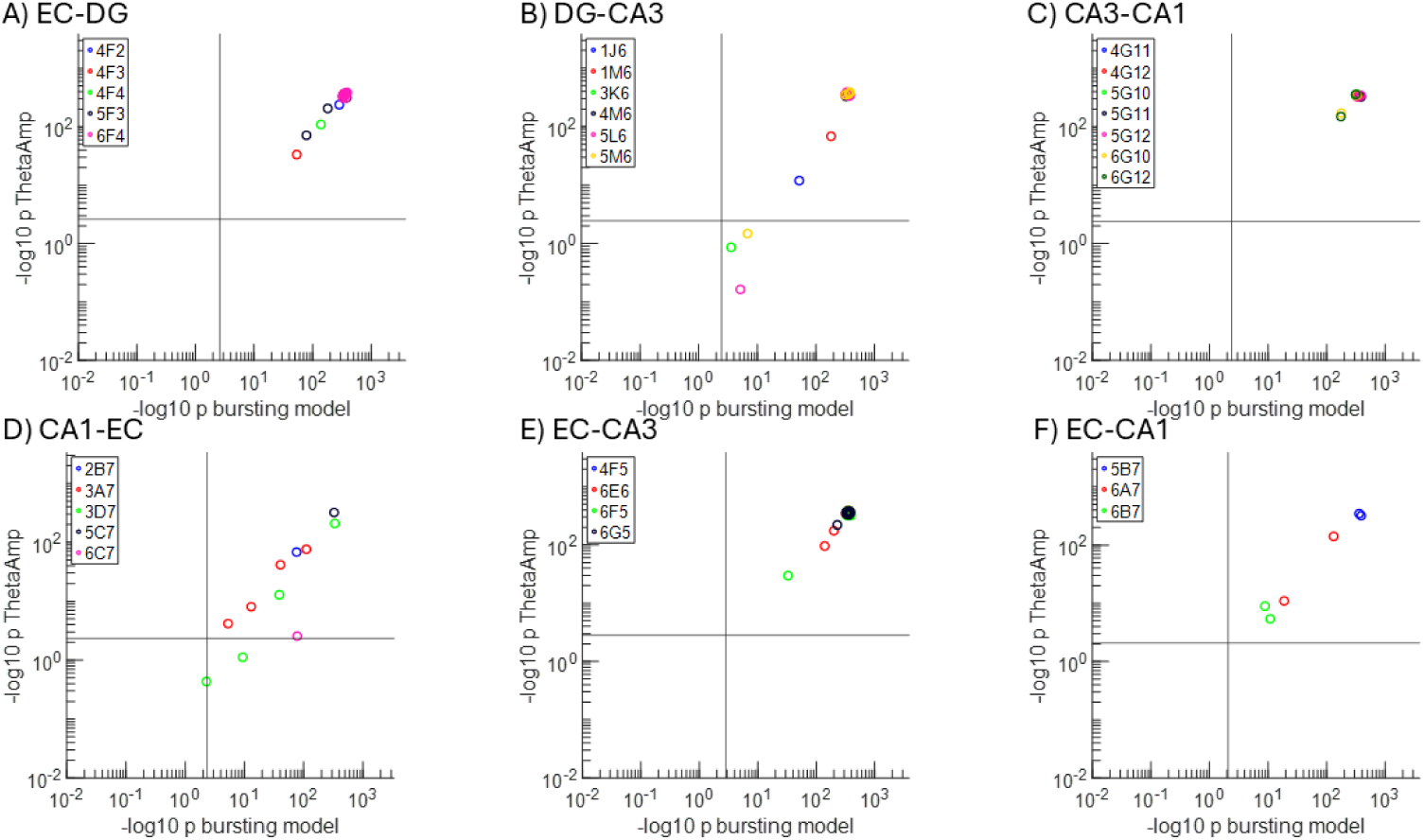
Many target neurons reliably respond with bursting associated with input axonal theta amplitude with the following frequencies: **A**) EC-DG 20 of 20 target somata, **B**) DG-CA3 11 of 14, **C**) CA3-CA1 12 of 12, **D**) CA1-EC 9 of 11, **E**) EC-CA3 34 of 34, and **F**) EC-CA1 6 of 6. Grey vertical and horizontal lines indicate the Bonferroni adjusted cutoffs for significance.

**Supplementary Fig. 8.**
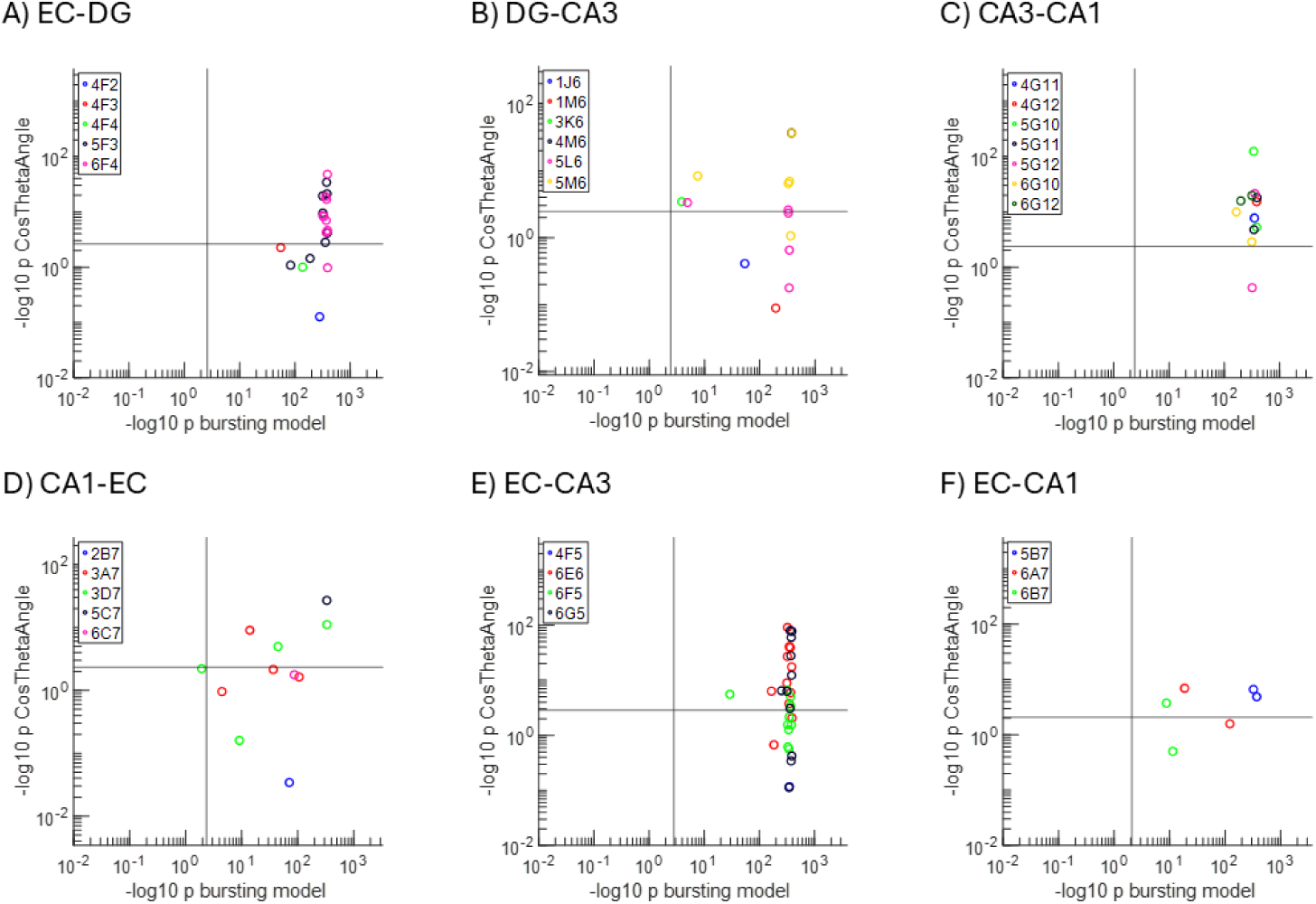
Many target neurons reliably respond with bursting associated with input axonal theta phase with the following frequencies: **A**) EC-DG 13 of 20, **B**) DG-CA3 8 of 14, **C**) CA3-CA1 11 of 12, **D**) CA1-EC 4 of 11, **E**) EC-CA3 21 of 34, and **F**) EC-CA1 4 of 6. Some p-values were measured at 0, so an epsilon to the 20th power was added to all numbers before the log transformation. A 5% jitter was added for graphical clarity. Grey vertical and horizontal lines indicate the Bonferroni adjusted cutoffs for significance. Some overlap of points at the extreme was due to machine precision limits.

### Rare coincident theta waves on two axons via mutual information

We tested the opposite of specificity for the occurrence of theta waves in a large fraction of axons by assessing their mutual information. For each feed-forward CA3-CA1axonal theta wave, Fig. **9**A-F shows very low coincident theta waves on all but one pair of CA3-CA1 axons in six networks. **Fig. 9G** shows the example of the one pair of high mutual information axonal theta oscillations, likely from a bifurcated axon. **Fig. 9H** one pair of medium mutual information axonal theta oscillations, possibly originating from two daughter neurons that shared a common input. This indicates that the same theta waves were rarely detected on two axons, suggesting that routing was unique on a per-unit basis. Other subregions shown in **Suppl Fig. 9** show all mutual information between feed-forward axons to be less than one bit, again showing low similarity in information between communicating axons.

**Fig. 9.**
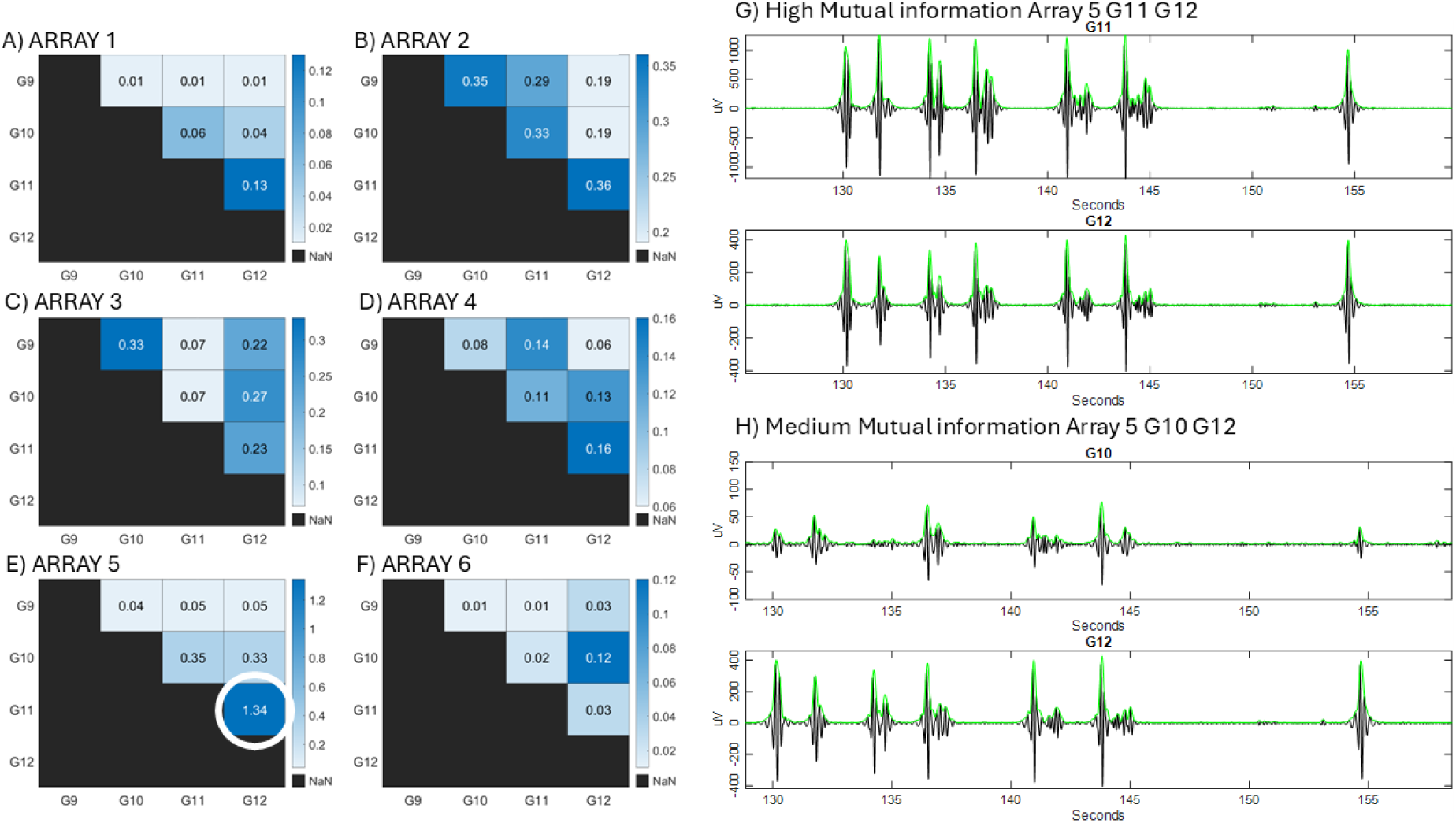
Non-redundancy of all 30 CA3-CA1 axons from six arrays; only one pair carried similar theta waves by mutual information. X and Y axes are electrodes over which axons pass in tunnels. Numbers are bits of mutual information for each electrode pair. **A-F**) Mutual information on each array compared with all other tunnels in CA3-CA1. **G**) Example of high mutual information (1.34) between axons G11 and G12 from FID 5 filtered for theta. **H**) Example of medium mutual information (0.35) between axons G10 and G12 filtered for theta show more variability in wave forms and amplitudes in G10.

**Supplementary Fig. 9.**
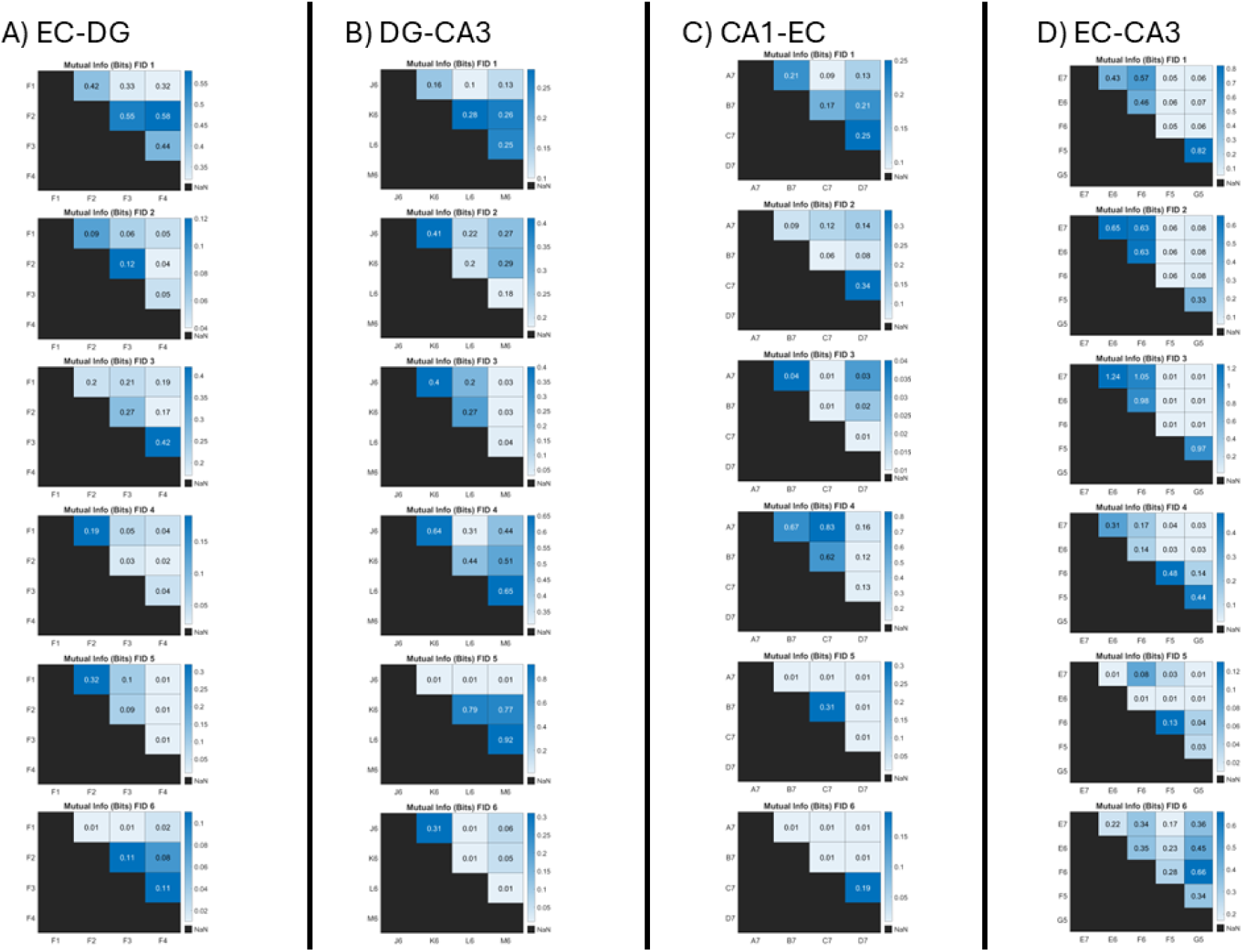
Mutual information of all inter-regions. **A-D**) Mutual information between axon pairs spanning the indicated subregions. Only FID 3 EC-CA3 had any mutual information bits greater than 1, however no targets were found for these axons in this array. All other tunnels had low mutual information.

## Discussion

We provided data to support the proposition that axons not only carry action potentials but also transmit subthreshold theta frequency voltage oscillations. Independent optogenetic inactivation of the EC or CA3 in live behaving rats showed that spiking and theta waves could be dissociated (Zutshi et al., 2022), as we show directly here in isolated axons. Although not proven, such a multiplex signal transmission satisfies some of the criteria that distinguish axonal theta waves from volume conduction and the synaptic currents. According to Box 1, these criteria include 1) sparsity, 2) theta voltage oscillations that originate at specific source somata, and 3) specific target neurons. If theta voltage oscillations are important for information transfer, then 4) responses will depend on the phase and amplitude of the theta wave. Further evidence is needed to show regenerative axonal subthreshold voltage oscillations for long distance transmission suggested by our inter-regional axonal transmission of theta waves.

### Sparsity of axons carrying theta frequency voltage oscillations between subregions

For a role of axonal theta in specific information transmission between subregions distinct from volume conduction, only a limited set of individual axons would carry theta waves and these would be different in different axons. This is sparse coding in a limited set of axons from the larger population, as shown for spatial learning in models of the mammalian visual cortex^31^. In our paper, we identified two features suggesting that the observed theta waves play a role in sparse encoding. First, we categorized the prevalence of high amplitude axonal theta in communicating axons. Out of all 21 tunnels monitored between subregions per array, single axons were identified in about 50% of monitored tunnels, the number of axons in a single inter-region ranged from 17% (2/12) to 27% (4/15) of axons with high amounts of theta. Second, by mutual information between communicating axons, we found that only two pairs of axons had mutual information above 1 bit out of 65 monitored single axon tunnels in five inter-regional sets from six arrays. This indicates that our model network satisfies the criteria for sparse encoding for axonal theta transmission between subregions and are not bathed in the same volume-conducted electric field and do not simply relay the same signals. In an artificial neural network, faster learning in densely connected networks was facilitated by sparse connectivity of information distributed across more nodes^32^. Graph theory analysis of our hippocampal networks also showed a balance of local (subregional) clustering with more limited inter-regional (sparse) connections (Lassers et al, 2023). The sparsity observed in our biological neural networks reflects an organizational pattern that balances dense local clustering with sparse long-range connections, thereby optimizing metabolic efficiency by reducing overall costs. Thus, sparse coding of theta oscillations in axons provides for greater statistical independence of output information.

### Identification of specific source neurons from which axonal theta waves originate

Identifying a source neuron from a limited set of inter-regional axons is a daunting task given the 11,000 estimated neurons in our model CA3 and only 18 subregion electrodes. Most of the neurons in a subregion would be connected locally, with only a few connections that reach far distances^33^. In the brain, this gives rise to small-world clusters of neurons that are responsible for local computation within subregions and few communicating axons between subregions. In our in vitro model, we looked for axonal theta waves whose amplitudes correlated with the axonal theta amplitudes in upstream subregions to identify the source. Out of 24 CA3-CA1 tunnels monitored from 6 arrays, only 3 axons had strong theta. For these 3 axons, we were able to find the source neuron or a neuron closely connected to the source in only 2 cases for CA3-CA1. This suggests that volume conduction of an electric field (LFP) could not be responsible for such a low percentage of source neuronal somata with strong theta coupled to a single axon.

### Identification of specific target neurons affected by axonal theta waves

Often, local field potentials are studied as the superposition of several dendritic postsynaptic potentials firing in synchrony^1, 34^. One assumed explanation, but not yet shown, is that this synchrony is due to a common afferent axon. In our work, we have identified several of these single axons with strong theta waves able to transmit these waves from one subregion to another AND alter the firing of specific target neurons. In vivo experiments that examine the coding of phase procession in the CA1 corroborate the CA3-CA1 axonal theta phase contribution to subregional spiking with phase centered around 0 degrees where the derivative of the voltage was greatest^35^, as we observed. In our experiments, the null hypothesis proposes that theta waves are unrelated to subregional spiking and lack impact on neuronal computation. Out of 65 total axons monitored, spiking in 38 target neurons were significantly modulated by strong theta in 56 targets with spiking out of 432 total subregional electrodes. If the axons were transmitting theta waves via volume conduction, many neurons in the subregion would be affected. Therefore, we reject the null hypothesis and say that transmission of theta phase and amplitude affects subregional target spiking and requires specificity of connections. While LFPs may be measured as a voltage spreading through tissue from the contribution of several synapses working in synchrony, we show that axons possibly transmit these theta frequency voltage oscillations to the dendrites. We next turn to how the targets are affected by the axonal theta waves.

### Dependence of target neuron spiking and bursting on phase and amplitude of the axonal theta voltage oscillations

We have discovered not just the existence of a direct connection between communicating axons and target subregions, but we have also described the modulation of target neuron activity by the axonal theta amplitude and phase. Neural systems often operate around log-log power-law dynamics, leading to a network state near a critical point for optimal information processing^15, 25, 36, 37^. These long-tailed linear distributions in the brain’s computations fit linear relationships in log-space and thus compute over a wider dynamic range than simple rate coding. Logarithmic encoding also enables multiplicative non-linear inputs to become additive^38^. We have found that the log of the axonal theta amplitude correlated strongly with the log of spike rate in certain target neurons. The phase of theta is also specific to the rate of subregional spiking. Over all of the axon-subregional target neuronal relationships, 4 of 6 excitatory pathways evoked the most spikes around zero degrees phase, at the rise of the voltage when the derivative of the phase is maximal for maximal current (I=CdV/dt), like those in the behaving rat^7^. The earlier arrival of the EC-CA3 axons to the CA3 target neuron synapses evoked minimal subregional spiking at this zero-degree phase angle may prime the target CA3 neurons for subsequent impact of DG theta. These qualitative relationships were evaluated quantitatively using the GLM of the Hilbert transform of theta amplitude and cosine of the phase angle to predict the probabilities of the target spiking and separately target bursting. The number of target neurons whose bursting was modulated by axonal theta was strongest for EC axons to the DG and CA3, supporting the oft cited EC as the source of theta wave generation in vivo^34, 39, 40, 41, 42^. This strong relationship for other axons in our model hippocampus may suggest independent generation of theta waves in each subregion. In vivo measures suggests an origin of theta from the frequency interaction of excitatory-inhibitory potentials^34^ and an important role in memory consolidation during human sleep^43^. Our work thus provides evidence for a subthreshold presynaptic membrane voltage oscillation that could drive theta frequency neurotransmitter release due to higher concentrations of calcium in the presynaptic terminal as hypothesized in the next section (**Fig. 10**).

**Fig. 10.**
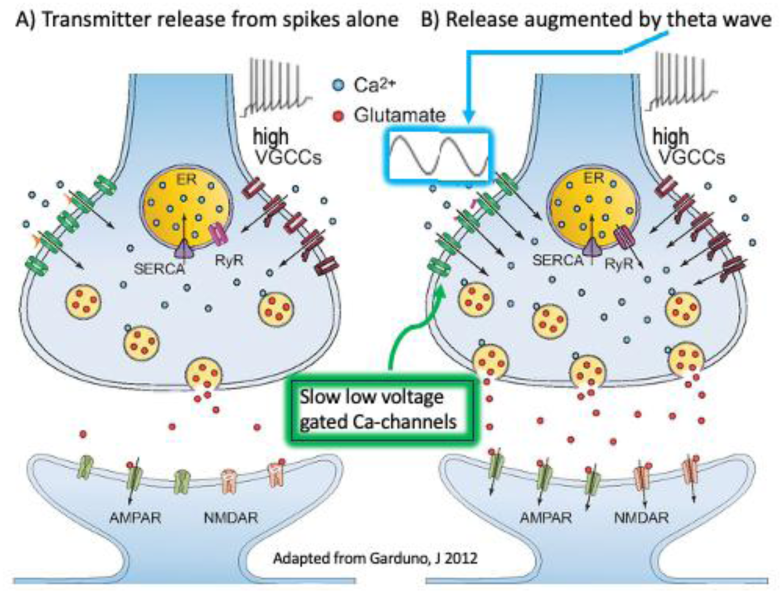
Compared to. **A**) a presynaptic terminal receiving only spikes, **B**) a terminal receiving a delta wave and spikes will activate more low voltage Ca-channels, admit more Ca^+2^ and release more transmitter (Adapted from Garduno 2012 J. Neurosci under CC-BY-NC-SA).

### Ion Channel Source of Theta Oscillations

The origin of local field potentials has been of great interest to the neuroscientific community for well over a decade^1^. Our work demonstrates that theta frequency power is not simply a byproduct of filtering spike amplitudes. Axonal subthreshold voltage oscillations at theta frequencies may involve slow low-voltage gated calcium channels, separate from Na^+^ and K^+^ ion channels of action potentials. At the presynaptic terminal, synaptic vessicle release is controlled by intracellular calcium levels. For axonal theta and other sub-threshold voltage oscillations to be transmitted through synapses, they must influence the release of synaptic vesicles. Computation and in vivo work have shown that CA1 pyramidal cells are specifically tuned to theta frequency input which affects threshold tuning for bursting activity^44, 45^. **Fig. 10** depicts a plausible mechanistic model mediated by slow-voltage gated calcium channels (VGCC) that increase calcium levels in presynaptic terminals to augment the well-known high VGCC activated by axonal action potentials and effect greater transmitter release. Candidate presynaptic low-voltage ion channels include Ca_v_3.1, 3.2, 3,3^46^. A second mechanistic requirement for axonal transmission of voltage oscillations at theta frequencies over long distances requires regenerative ion channel mechanisms. We have provided evidence that theta is regenerative due to long-distance transmission from identified source neuron, over axonal transmission ofat least 200 micrometers based on electrode spacing. We have found grandparent sources over 1.1 mm away that exists without volume conduction. However, we have not identified the specific ion channels responsible for the generation and transmission of theta. This will require further work and pharmacological intervention to identify and categorize the ion channels involved.

### Theta-waves and multiplex coding in spiking neural networks (SNN) models

Recent *in silico* neural network advancements in have attempted to switch from convolutional neural networks (CNNs) to spiking neural networks (SNNs) to take advantage of the inherent temporal encoding features of event based computing^47, 48, 49^. In SNNs, action potentials are simulated using a low-cost leaky integrate and fire (LIF) model that approximates the Hodgkin-Huxley model of biological action potentials^49^. However, these SNNs running on LIF neuron models stop at the simulation of spikes and do not include the oscillitory contributions of local fields. We have shown in this paper that LFPs are separate phenomena from spiking but contribute to the synchronization of spikes across long distances through amplitude and phase dynamics. These subthreshold theta dynamics as found in our work can be applied to current SNNs to move the models closer to biological representations. The first attempts at simulating subthreshold oscillations was with resonate-and-fire neurons^50^. By representing the state variable in the LIF neuron as complex, pulses could cause the voltage decay to act as a dampened oscillator. The resonate-and-fire model was improved through the linearization of the FitzHugh-Nagumo neural model near an hyberbolic fixed point and showed that phase locking can occur with pertubations from spikes even in the presence of noise. Furthermore, incorporation of oscillitory signals in addition to standard leaky integrate and fire exponential decays greatly improved performance of event-based benchmark datasets^51^. Incorporating heterogeneous subthreshold oscillations into SNNs produced more energy efficient and noise-resistant temporal processing^52^. We propose that our results can be used to inform future SNN models on the specific parameters to incorporate into the network’s oscillatory behavior.

### Limitations and relevance to the in vivo brain

The most pressing concern of our experimental setup must be its relevance to *in vivo* brain behavior and physiology. Hundreds of papers have used theta burst stimulation of the perforant path to induce LTP in hippocampal slices^9, 53, 54^. This in situ approach could be generating the very axonal theta frequency subthreshold waves that we observed directly. Further, adjustment of the amplitude of the theta burst currents used in slices to achieve a faster or stronger population burst in CA1 may relate to the axonal theta amplitude effects on target spiking that we report. In another area, early studies showed that lesion of septal inputs eliminated theta oscillations essential for spatial memory^40^. We found theta oscillations without modulatory cholinergic septal inputs, which argues against septal genesis of axonal theta waves. In addition to cholinergic modulation contributing to theta, NMDA and GABAA/B contribute to theta in slices^55^. In the DG of freely behaving rats, LTP was enhanced by stimulation in phase with theta oscillations and inhibited by antiphase stimulation^56^. Sandler (2015) found in vivo that CA1 theta required a complete loop of the circuit, suggesting a need for axonal transmission. Hoppenstead and Izhikevich (2000) have demonstrated theoretical precedent of phase encoding in phase locked loops. Local field potentials have been associated with the consolidation of memory during sleep, and have been associated with up and down states of activity in human brain slices^43^, which clearly requires thalamic inputs that we lack. Yet our model still has intrinsic mechanisms for axonal theta generation.

We realize our in vitro model is a two-dimensional network which lowers the degrees of freedom for network complexity from the 3D brain. While this increases the probability of finding the direct connections between source neuron, axon and target neuron, it can’t represent the laminated arrangements of neurons seen in the mammalian hippocampus and certainly lacks important modulatory inputs from other brain regions and perihippocampal and neocortical inputs present that would typically integrate information at the EC. Our previous work has pushed us toward creating a more biologically accurate model of the trisynaptic loop by including the EC-CA3 perforant pathway^17^ which expanded on previous analysis. With the addition of this pathway, we can see the physiological fan-out as most number of targets in the EC-DG and EC-CA3 paths that are expected *in vivo* and more robust bursting behavior in axons from CA3 to CA1. In vivo models found the strongest theta power was generated in the EC in behaving rats^39^, from which we saw routing to the most prevalent DG and CA3 targets in our model. Similar to the GLM relationships we see between axonal theta and subregional target spiking in multiple subregions, in vivo studies have shown theta phase relationships to the gamma band associated with individual spiking activity in multiple hippocampal subregions^57^. While our simple network complexity only crudely reflects the biological organization of the *in vivo* hippocampus, we may have gained insight into axonal communication and dynamics between subregions. Future directions of this work will examine the coherence of theta and gamma waves in axons and how axonal gamma coordinates spiking activity in subregions. Identification of the ion channels responsible for regenerative propagation of subthreshold oscillations will reveal mechanistic targets for manipulating network activity.

## Conclusion

We have presented evidence that an in vitro model hippocampal network produces sparse, specific theta waves isolated to single axons that affects subregional spiking and bursting based on theta amplitude and phase angle. Beyond the existence of LFPs that arrise from the summed EPSPs in synchronously active neurons^1^, the regenerative axonal transmission of subthreshold transmembrane voltage oscillations better explains synchronization of spiking and LFP activity between subregions. This paradigm shift of axonal transmission of theta waves better explains sparse interregional communication than the 1/r^2^ fall off via volume conduction of voltages from synaptic current sources^1, 6^. These findings better illuminate the mesoscopic mechanisms and dynamics for spiking and bursting coordination between subregions of the hippocampus. Based on these findings, we can now search for the ionic mechanisms at play that are responsible for generating subthreshold oscillations. Our finding of axonal theta waves helps to explain the communication through coherence hypothesis^58^ that enables phase-locked communication between distant subregions.

## Acknowledgements

This work was supported in part by funds from the UC Irvine Foundation. We would also like to thank Drs. György Buzsáki, Michael Hasselmo, and Jennifer Gelinas for their expertise in this field and guiding comments while preparing the manuscript.

## Author information

### Authors and Affiliations

1) Department of Biomedical Engineering, University of California, Irvine, CA, 92697, USA Sam Lassers, Gregory Brewer
2) Center for Neurobiology of Learning and Memory and MIND Center, University of California, Irvine, CA, 92697, USA Gregory Brewer

## Contributions

SBL wrote the scripts to analyze axonal direction, calculate theta wave modulation index and mutual information, and compute general linear models for theta amplitude and phase angles predicting spiking. GJB funded and conceptualized the experiments and recorded the original data. Both SBL and GJB wrote all drafts of the manuscript.

## Corresponding author

Correspondence to Gregory Brewer.

## Ethics declarations

### Animal use protocol

The animal study was approved by UC Irvine Institutional Animal Care and Use Committee under AUC 23-013. The study was conducted in accordance with the local legislation and institutional requirements.

## Data Availability

The data used in this study is freely available on Data Dryad. The code is available in our [GitHub repository](https://github.com/Brewer-Neurolab/Lassers_Spike_LFP).

